# Quantifying GC-biased gene conversion in great ape genomes using polymorphism-aware models

**DOI:** 10.1101/380246

**Authors:** Rui Borges, Gergely Szöllősi, Carolin Kosiol

## Abstract

As multi-individual population-scale data is becoming available, more-complex modeling strategies are needed to quantify the genome-wide patterns of nucleotide usage and associated mechanisms of evolution. Recently, the multivariate neutral Moran model was proposed. However, it was shown insufficient to explain the distribution of alleles in great apes. Here, we propose a new model that includes allelic selection. Our theoretical results constitute the basis of a new Bayesian framework to estimate mutation rates and selection coefficients from population data. We employ the new framework to a great ape dataset at we found patterns of allelic selection that match those of genome-wide GC-biased gene conversion (gBCG). In particular, we show that great apes have patterns of allelic selection that vary in intensity, a feature that we correlated with the great apes’ distinct demographies. We also demonstrate that the AT/GC toggling effect decreases the probability of a substitution, promoting more polymorphisms in the base composition of great ape genomes. We further assess the impact of CG-bias in molecular analysis and we find that mutation rates and genetic distances are estimated under bias when gBGC is not properly accounted. Our results contribute to the discussion on the tempo and mode of gBGC evolution, while stressing the need for gBGC-aware models in population genetics and phylogenetics.

## 1 Introduction

The field of molecular population genetics is currently been revolutionized by progress in data acquisition. New challenges are emerging as new lines of inquiry are posed by increasingly large population-scale sequence data (Casillas and Barbadilla, 2017). Mathematical theory describing population dynamics has been developed before molecular sequences were available (e.g. Fisher (1930); Wright (1931); Moran (1958); Kimura (1964)); now that ample data is available to perform statistical inference, many models have been revisited. Recently the multivariate Moran model with boundary mutations was developed and applied to exome-wide allele frequency data from great ape populations (Schrempf and Hobolth, 2017). However, drift and mutation are not fully sufficient to explain the observed allele counts (Schrempf and Hobolth, 2017). It was hypothesized that other forces, such as directional selection and GC-biased gene conversion (gBGC), may also play a role in shaping the distribution of alleles in great apes.

Directional selection and gBGC have different causes but similar signatures: under directional selection, the advantageous allele increases as a consequence of differences in survival and reproduction among different phenotypes; under gBGC, the GC alleles are systematically preferred. gBGC is a recombination-associated segregation bias that favors GC-alleles (or strong alleles, hereafter) over AT-alleles (or weak alleles, hereafter) during the repair of mismatches that occur within heteroduplex DNA during meiotic recombination (Marais, 2003). gBGC was studied in several taxa including mammals (Duret and Galtier, 2009; Romiguier et al., 2010; Lartillot, 2013), birds (Webster et al., 2006; Weber et al., 2014; Sme, 2016; Corcoran et al., 2017), reptiles (Figuet et al., 2015), plants (Muyle et al., 2011; Liu, 2018; Clément et al., 2017; Serres-Giardi et al., 2012) and fungi (Pessia et al., 2012; Lesecque et al., 2013; Liu, 2018). However, apart from some studies in human populations (Katzman et al., 2011; Glémin et al., 2015; Pouyet et al., 2018), a population-level perspective of the intensity and diversity of patterns of gBGC among closely related populations is still lacking.

Several questions remain open regarding the tempo and mode of gBGC evolution. For example, the effect of demography on gBGC is still controversial. While theoretical results and studies in mammals and birds advocate a positive relationship between the effective population size and the intensity of gBGC (Nagylaki, 1983; Romiguier et al., 2010; Weber et al., 2014), Galtier et al. (2018) failed to detect such relationship between animal phyla. These results open the question to which extent demography shapes the intensity of gBGC in closely- versus distant-related species/populations. Another aspect that is not completely understood is the impact of GC-bias on the base composition of genomes (Phillips et al., 2004; Romiguier et al., 2013). In particular, the individual and joint effect of gBGC and mutations shaping the substitution process remains elusive. Here, we address these two questions by revisiting the great ape data (Prado-Martinez et al., 2013) with a Moran model that accounts for allelic selection.

The Moran model (Moran, 1958) has a central position describing populations’ evolution: it models the dynamics of allele frequency changes in a finite haploid population. Recently, an approximate solution for the multivariate Moran model with boundary mutations (i.e. low mutation rates) was derived (Schrempf and Hobolth, 2017). In particular, the stationary distribution was shown useful to infer population parameters from allele frequency data (Schrempf et al., 2016; Schrempf and Hobolth, 2017). Here, we present the Moran model with boundary mutations and allelic selection, derive the stationary distribution, and we build a Bayesian framework to estimate population parameters.

Other approaches making use of allele frequency data to estimate mutation rates and selection coefficients have been proposed in the literature. Glémin et al. (2015) proposed a method to quantify gBGC from the derived allele frequency spectra that incorporates polarization errors, takes spatial heterogeneity into account, and jointly estimates mutation bias. The number of derived alleles is modelled by a Poisson distribution on the mutation rates among weak, strong a neutral alleles (Muyle et al., 2011). Our approach differs from Glémin et al. (2015) as it does not require polarized data or need to account for polarization errors. In addition, our method makes use of the information given by the fixed sites, information that is usually discarded by other methods (Glémin et al. (2015) included).

Furthermore, our application to great apes shows that most great apes have patterns of allelic selection consistent with gBGC. Our results suggest further that demography has a major role in determining the intensity of gBGC among great apes, as the intensity of the obtained selection coefficients significantly correlates with the effective population size of great apes. We also show that not accounting for GC-bias may considerably distort the reconstructed evolutionary process, as mutation and substitution rates are estimated under bias.

## 2 Methods

### 2.1 The multivariate Moran model with allelic selection

The modelling framework defined in this work builds in the model described by Schrempf et al. (2016), which according to some proposed terminology (Vogl and Bergman, 2015; Schrempf and Hobolth, 2017), can be addressed as the multivariate Moran model with boundary mutations (hereafter *MM*). Here, we described the multivariate Moran model with boundary mutations and allelic selection (hereafter *MS*). The multivariate Moran model can be also referred as a polymorphism-aware phylogenetic model (PoMo) if we consider 4-variate case (De Maio et al., 2013, 2015; Schrempf et al., 2016), those representing the 4 nucleotide bases (Figure 1).

**Figure 1:**
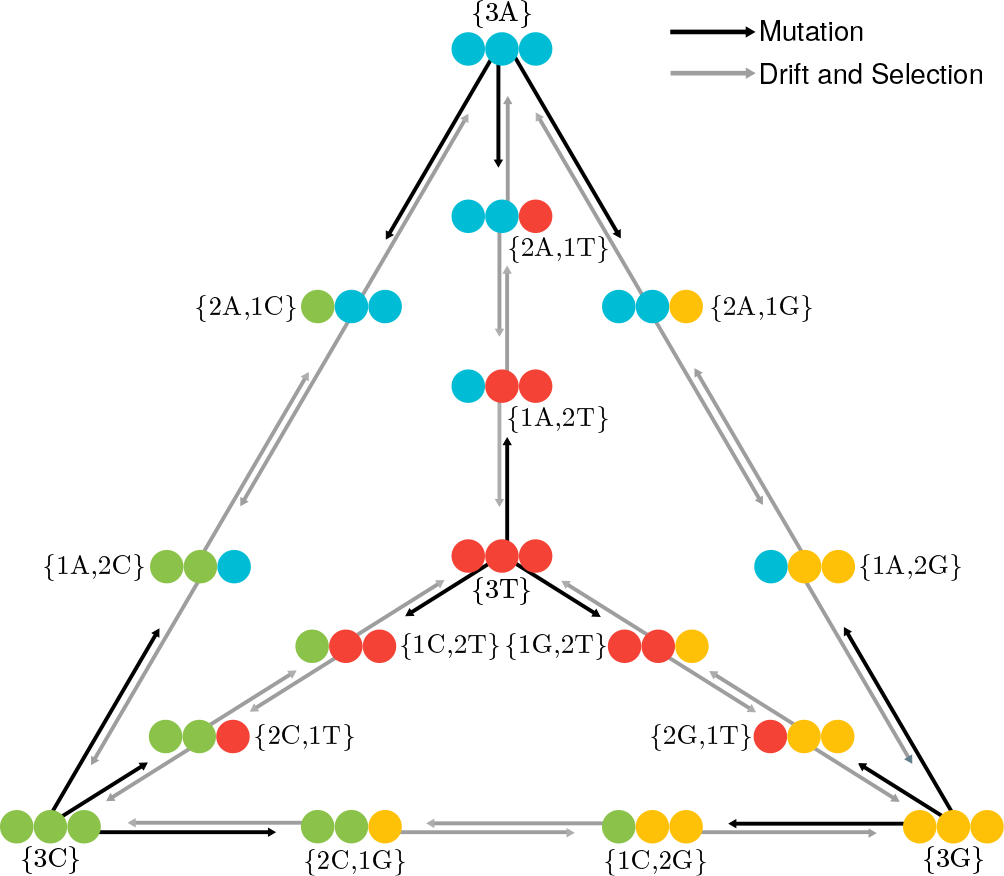
PoMo state-space using *N* = 3. The 4 alleles represent the four nucleotide bases. Black and grey arrows indicate mutations, and genetic drift plus selection, respectively. Monomorphic or boundary states {*Na*_*i*_} are represented in the tetrahedron’s vertices, while the polyMorphic states {*na*_*i*_ (*N−n*)*a*_*j*_} are represented in its edges. Monomorphic states interact with polymorphic states via mutation, but a polymorphic state can only reach a monomorphic state via drift or selection. Between polymorphic states only drift and selection events occur.

Consider a haploid population of *N* individuals and a single locus with *K* alleles: *a*_*i*_ and *a*_*j*_ are two possible alleles. The evolution of this population in the course of time is described by a continuous-time Markov chain with a discrete character-space defined by *N* and *K*, each of which represents a specific assortment of alleles. Two types of states can be defined: if all the individuals in a populations have the same allele, the population is monomorphic {*Na*_*i*_}, i.e. the *N* individuals have the allele *a*_*i*_; differently, if two alleles are present in the population, the population is polymorphic {*na*_*i*_, (*N−n*)*a*_*j*_}, meaning that *n* individuals have the allele *a*_*i*_ and (*N − n*) have the allele *a*_*j*_. *n* is therefore the absolute frequency of allele *a*_*i*_ in the population.

Alleles trajectories are given by the rate matrix ***Q***. Time is accelerated by a factor of *N*, and therefore instead of describing the Moran dynamics in terms of Moran events (Moran, 1958), we developed a continuous version in which the time is measured in number of generations.

Drift is defined by the neutral Moran model: the transition rates of the allelic frequency shifts, only depend on the allele frequency and are therefore equal regardless the allele increases or decreases in the population (Durrett, 2008)

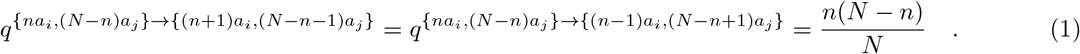

We accommodated mutation and selection in the framework of the neutral Moran model by assuming them to be decoupled (Baake and Bialowons, 2008; Etheridge et al., 2010).

Mutation is incorporated based on a boundary mutation model, in which mutations only occur in the boundary states. The boundary mutations assumption is met if the mutation rates 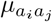 are small (and *N* is not too large). More specifically, Schrempf et al. (2016) established that 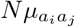 should be lower than 0.1, by comparing the expectations of the diffusion equation with the polymorphic diversity under the Moran model. In fact, most eukaryotes fulfill this condition (see Lynch et al. (2016) for a review of mutation rates). Another assumption of our boundary mutation model is that the polymorphic states can only be biallelic. However, this assumption is not a significant constraint as tri-or-more allelic sites are rare in sequences with low mutation rates.

We employed the strategy used by Burden and Tang (2016) and separated our model into a time-reversible and a flux part. We wrote the mutation rates as the entries of a specific mutation model, the general time-reversible model (GTR) (Tavaré, 1986): 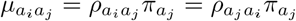, where ***ρ*** represents the exchangeabilities between any two alleles and ***π*** the allele base composition (equation (2)). Here, we restricted ourselves to the GTR, as this model simplifies obtaining formal results (Burden and Tang, 2016). Because ***π*** has *K* − 1 free parameters and ***ρ*** includes the exchangeabilities for all the possible pairwise combinations of *K* alleles, we ended up having *K*(*K* + 1)/2 − 1 free parameters in the GTR-based boundary mutation model.

Until now we have essentially described the model proposed by (Schrempf et al., 2016); this work extends this model by including allelic selection. We modeled allelic selection by defining *K* − 1 relative selection coefficients ***σ***: an arbitrary selection coefficient is fixed to 0. The selection coefficients defined this way is guaranteeing that our multi-allelic model behaves neutrally only under the condition that all the selection coefficients are the same and equal to 0. Defining the fitness as the probability that an offspring of allele *a*_*i*_ is replaced with probability 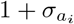 (Durrett, 2008), we can formulate the component of allelic selection alongside with drift, and thus among the polymorphic states (equation (2)).

Altogether, the instantaneous rate matrix ***Q*** of the multivariate Moran model with boundary mutations and allelic selection can be defined as

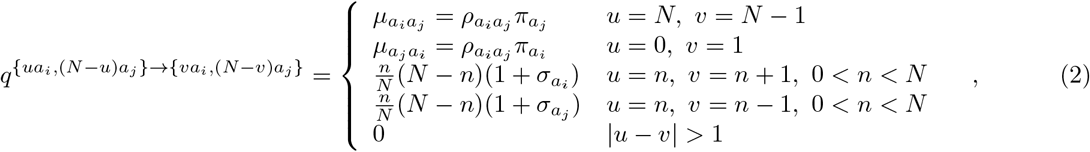

where *u* and *v* represent a frequency change in the allele counts (though *N* remains constant). The diagonal elements are defined by the mathematical requirement such that the respective row sum is 0.

As the parameters of the population size, mutation rate and selection coefficients are confined, it is possible to scale down them to a small value *N* while keeping the overall dynamics unchanged (appendix A). The virtual population size *N* becomes a parameter describing the number of bins the allele frequencies can fall into. As a result, we can think of *N* either as a population size or a discretization scheme.

### 2.2 The stationary distribution

The stationary distribution of a Markov process can be obtained by computing the vector ***ψ*** satisfying the condition ***ψQ*** = **0**(appendix B). ***ψ*** is the normalized stationary vector and has the solution

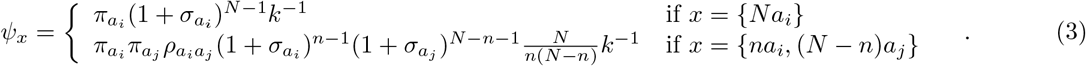

*k* is the normalization constant

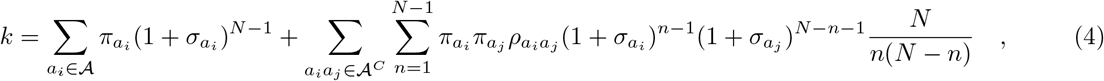

where 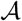 is the alphabet of the *K* alleles {*a*_1_, …, *a*_*K*_ }, representing the monomorphic states, and 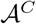 all the possible pairwise combinations of 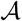 representing the *K*(*K* − 1)/2 types of polymorphic states {*a*_1_*a*_2_, *a*_1_*a*_3_, …, *a*_*K*−1_*a*_*K*_ }. For example, for the 4-multivariate case we write 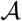 as the alphabet of the 4 nucleotide bases {*A, C, G, T*} and 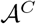 as all the possible pairwise combinations of the four nucleotide bases {*AC, AG, AT, CG, CT, GT*}. For a population of size *N* we have 4 + 6(*N* − 1) possible states, four of which are monomorphic (Figure 1).

### 2.3 Expected number of Moran events

From ***Q*** and ***ψ***, we can compute the expected number of Moran events (mutations, drift and selection) or the expected divergence per unit of time (in generations) under the *MS* model (appendix C):

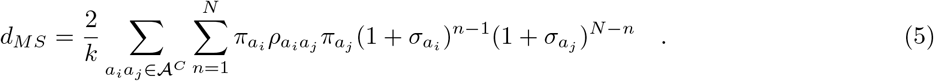

The quantity (5) can also be interpreted as the overall rate of the model. The expected number of Moran events for the neutral model can be easily calculated by letting ***σ*** → **0**. To compare the Moran distance *d*_*MS*_ with the standard models of evolution, we recalculated the Moran distance to only account for substitutions events 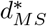: we corrected *d*_*MS*_ by the probability of a mutation and a subsequent fixation under the Moran model (appendix D)

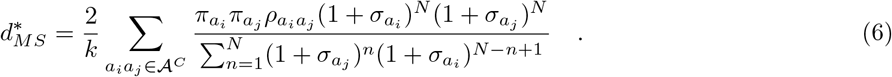

### 2.4 Bayesian inference with the stationary distribution

We can define a likelihood function based on the stationary distribution for a set of *S* independent sites in *N* individuals by taking the product of *ψ*_*x*_ over counts of monomorphic and polymorphic sites *c*(*x*)

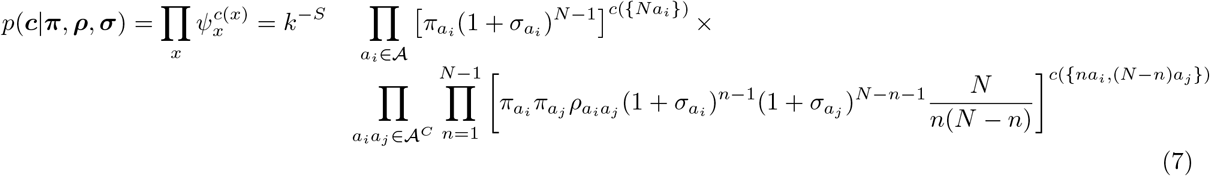

We employed a Bayesian approach: we define the prior distributions independently, a Dirichlet prior for ***π*** and an exponential prior for ***ρ*** and ***σ***; a Dirichlet and multiplier proposals were set for the aforementioned parameters with tuning parameters guaranteeing a target acceptance rate of 0.234 (Roberts et al., 1997). We employed the Metropolis-Hastings algorithm (Hastings, 1970) for each conditional posterior in a Markov chain Monte Carlo sequence to obtain random samples from the posterior. The algorithm was coded in the R statistical programing language (R Core Team, 2015): the packages `MCMCpack` and `expm` were integrated in our code to obtain samples from the Dirichlet density and to compute the matrix exponential, respectively (Martin et al., 2011; Goulet et al., 2017).

### 2.5 Application: great ape population data

The stationary distribution of 4-multivariate model was employed to infer the distribution of allele frequencies, selection coefficients and mutation rates from 4-fold degenerate sites of exome-wide population data from great apes (Prado-Martinez et al., 2013). We used 11 populations with up to 27 diploid individuals, totaling ~2.8 million sites per population (Table 1). Data preparation follows the pipeline described in De Maio et al. (2015). Estimates of the Watterson’s *θ* genetic diversity is below 0.003 for all the studied populations (Schrempf et al., 2016), validating the boundary mutations assumption of 0.1.

### 2.6 Data availability

Data and R scripts necessary to confirm the findings of this article are available on the GitHub (https://github.com/pomo-dev/pomo_selection). Supplemental material is available on Figshare and bioRxiv (https://doi.org/10.1101/380246).

## 3 Results

### 3.1 Simulations and algorithm validation

To validate the analytical solution for the stationary distribution of the multivariate Moran model, we compare it to the numerical solution obtained by calculating the probability matrix of ***Q**t* for large enough *t*. We confirmed that the numerical solution converges to the analytical solution (Figure S1).

We validated the Bayesian algorithm for estimating population parameters from the stationary distribution by performing simulations (Table S2 and Figures S2-S5). Our algorithm efficiently recovers the true population parameters from simulated allele counts. We tested the algorithms for different number of sites (10^3^, 10^6^ and 10^9^) and state-spaces (*N* = 5, 10 and 50). The number of sites does not increase the computation time substantially and is not a limiting factor for genome-wide analysis. In contrast, the size of the state-space influences the computational time. For larger state-spaces *N*, more iterations are needed to obtain converged, independent and mixed MCMC chains during the posterior estimation.

### 3.2 Patterns of allelic selection in great apes

To test the role of allelic selection defining the distribution of alleles in the great apes, we compared the neutral multivariate Moran model (*MM*) and the model with allelic selection (*MS*). Using the predictive stationary distribution and the observed allele counts, we computed the Bayes factors (BF) favoring the more complex model *MS* (i.e. log BF > 0 favors the model with allelic selection) for all populations. It is clear that *MS* fits the data considerably better for most of the studied great apes (log BF > 100, Table 1). The only exception is the Eastern gorillas population, for each a lower log BF was obtained (log BF = 5.497, Table 1).

**Table 1:**
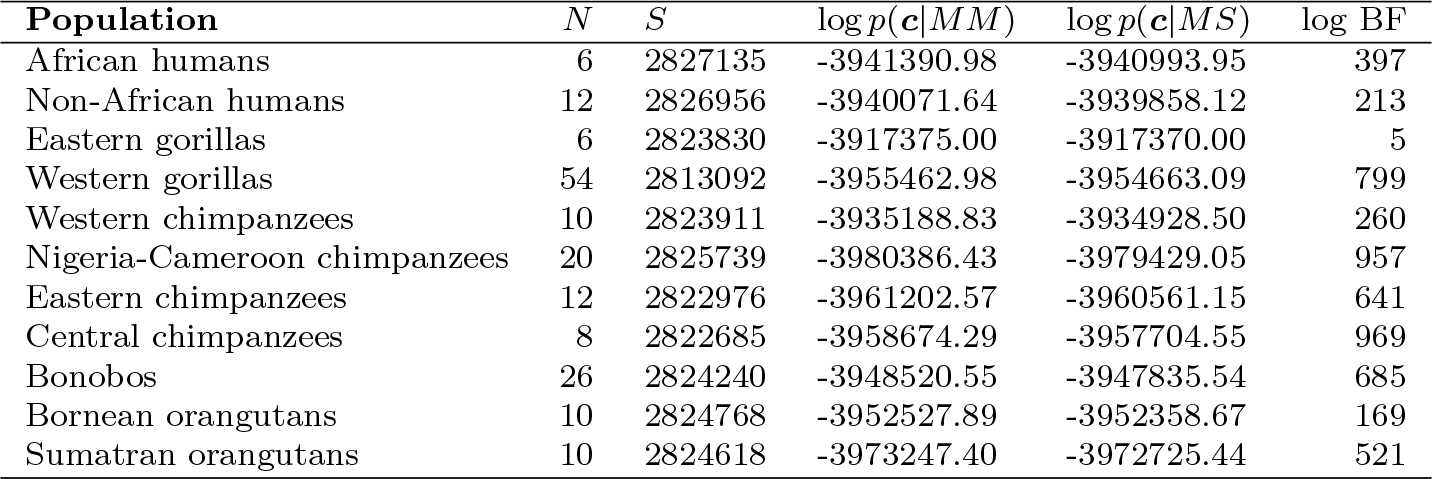
Evidence of allelic selection among the great ape populations. The number of haploid individuals and the number 4-fold degenerate sites per population are indicated by *N* and *S*, respectively. The log Bayes factors (log BF) were calculated as the sum over the product of the allele counts ***c*** and the posterior predictive probabilities under the Moran model with boundary mutations (*MM*) and allelic selection (*MS*). BF favor the model with allelic selection when higher than 1.

We have also corroborated our BF by inspecting the fit of the predictive distribution of *MM* and *MS* with the allele counts (Figure S6A-K). The allele counts for the polymorphic states are not symmetric, generally one allele if preferred and so are the polymorphic states that have it in higher proportions. As expected, we observed that *MS* better reproduces the skewed distribution of allele counts among great apes.

We further investigated the patterns of allelic selection in great apes by analyzing the posterior distribution of the relative selection coefficients of C, G, and T (*σ*_*A*_ was set to 0) under *MS*. A general pattern of allelic selection is observed in great apes: the selection coefficients of C and G are similar (meaning that their posterior distributions largely overlap), but different from the selection coefficient of T, which in turn overlaps 0 (approximately equal to the selection coefficient of A) (Figure 2). The only exception is the Eastern gorillas, for which the selection coefficients are all only slightly higher than 0 and rather similar (Figure 2). This result corroborates the relatively low BF found for evidence of allelic selection in the Eastern gorilla population.

**Figure 2:**
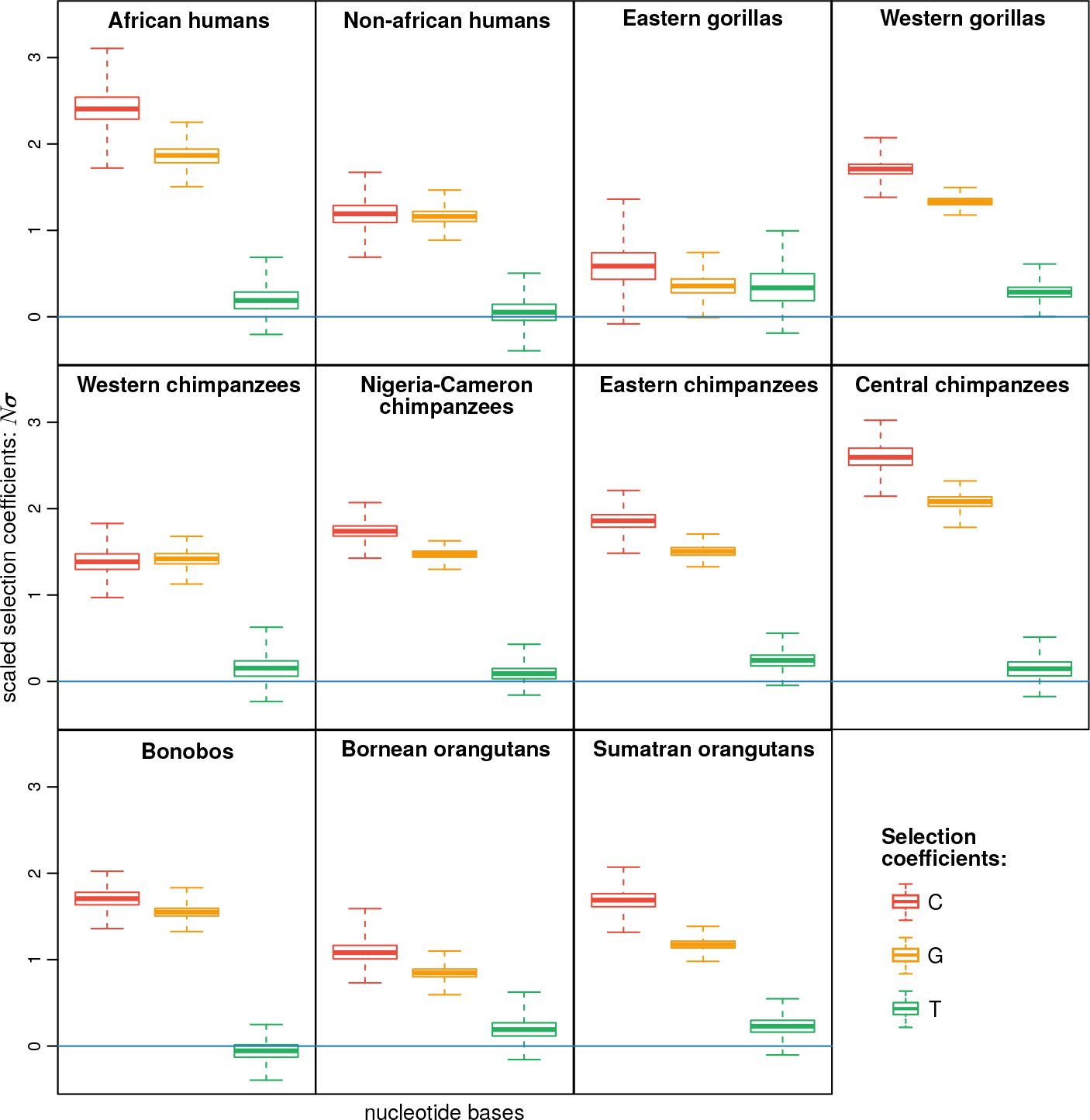
Scaled allelic selection coefficients for the great apes 4-fold degenerate synonymous sites. The boxplots represent the posterior distribution of the C, G and T scaled selection coefficients (*σ*_*A*_ was set to 0); the estimates were obtained using the 4-variate Moran model. The line in blue represents *σ*_*A*_ = 0. Table S3 summarizes the average scaled selection coefficients for each great ape population.

We further explored this result in order to check if the patterns of GC-bias found among great apes can be associated with gBGC. We correlate the GC-bias per chromosome (*σ*_*C*_ + *σ*_*G*_) with the chromosome size and recombination rate in the non-African human population (Figure S7), for which this data is particularly well characterized (Jensen-Seaman, 2004). We found a significant positive correlation between the GC-bias and recombination rate (Spearman’s *ρ* = 0.57, *p*-value = 0.006), but a negative correlation with the chromosome length (Spearman’s *ρ* = −0.52, *p*-value = 0.014).

Although the patterns of selection among great apes are similar qualitatively, they differ quantitatively. For example, the Central chimpanzees have patterns of GC-bias around 2.08/2.60 (*σ*_*C*_/*σ*_*G*_, Table S3 and Figure 2), while the closely related population of Western chimpanzees shows less strong patterns (around 1.38/1.42). Likewise, the GC-bias content in African and non-African human populations contrasts: 2.41/1.86 and 1.19/1.16, respectively. These results show that the patterns of allelic selection greatly vary among great apes, even among closely related populations.

It has been hypothesized that gBGC is a compensation mechanism for the mutational bias that exists in favor of the weak alleles, A and T (Duret and Galtier, 2009; Philippe et al., 2011): the AT/GC toggling effect. We observed that mutation rates from strong to weak alleles are more frequent (by a factor of 2.80; 3.26 if the stationary frequencies are accounted), while no mutational bias was found between alleles of the same type (1.02; 0.98 if the stationary frequencies are accounted; supplementary table S3). As the estimated selection coefficients have a clear pattern of GC-bias in most great apes, we can conclude that our analyses are congruent with the expectations of the AT/GC toggling effect.

Furthermore, we compared our method with Glémin et al. (2015), by considering only two alleles (the strong (S) and weak (W) alleles) using human allele counts from the first human chromosome, divided into 51 regions of 1 million sites (data taken from Glémin et al. (2015)). We compared estimates of the gBGC rate coefficient as predicted by our model and that of Glémin et al. (2015) (*σ*_*S*_ and *B*, respectively) and observed that they are negatively correlated (Spearman’s *ρ* = −0.37, *p*−value = 0.012). Interestingly, *B* significantly correlates with our estimates of *µ*_*WS*_ (the mutation rate of weak to strong alleles; *ρ* = −0.50, *p−*value = 0.001). We have further checked the influence of the fixed sites in our estimates of gBGC and, as expected, we observed that *σ*_*S*_ positively correlates with the percentage of monomorphic sites (*ρ* = −0.36, *p−*value = 0.012); intriguingly, *B* is negatively correlated (*ρ* = −0.46, *p−*value = 0.001). Scatter plots of the mentioned correlation tests can all be found in figure S8.

### 3.3 *N*_*e*_ and the total rate of mutation and selection in great apes

It is widely known that the intensity of mutation and selection reflect population demography. To check whether the estimated mutation and selection coefficients among great ape populations may be explained by demography, we tested the correlation between the total rate of mutation and selection and *N*_*e*_ (obtained from Tenesa et al. (2007); Prado-Martinez et al. (2013)). Positive correlations between the total mutation and selection rates and the effective population size were obtained (Figure 3): Pearson’s correlation coefficient of 0.59 (*p*-value = 0.070) and 0.87 (*p*-value = 0.001), respectively. These correlations were obtained using independent contrasts (Felsenstein, 1985) accounting for the great apes phylogeny as predicted in Prado-Martinez et al. (2013).

**Figure 3:**
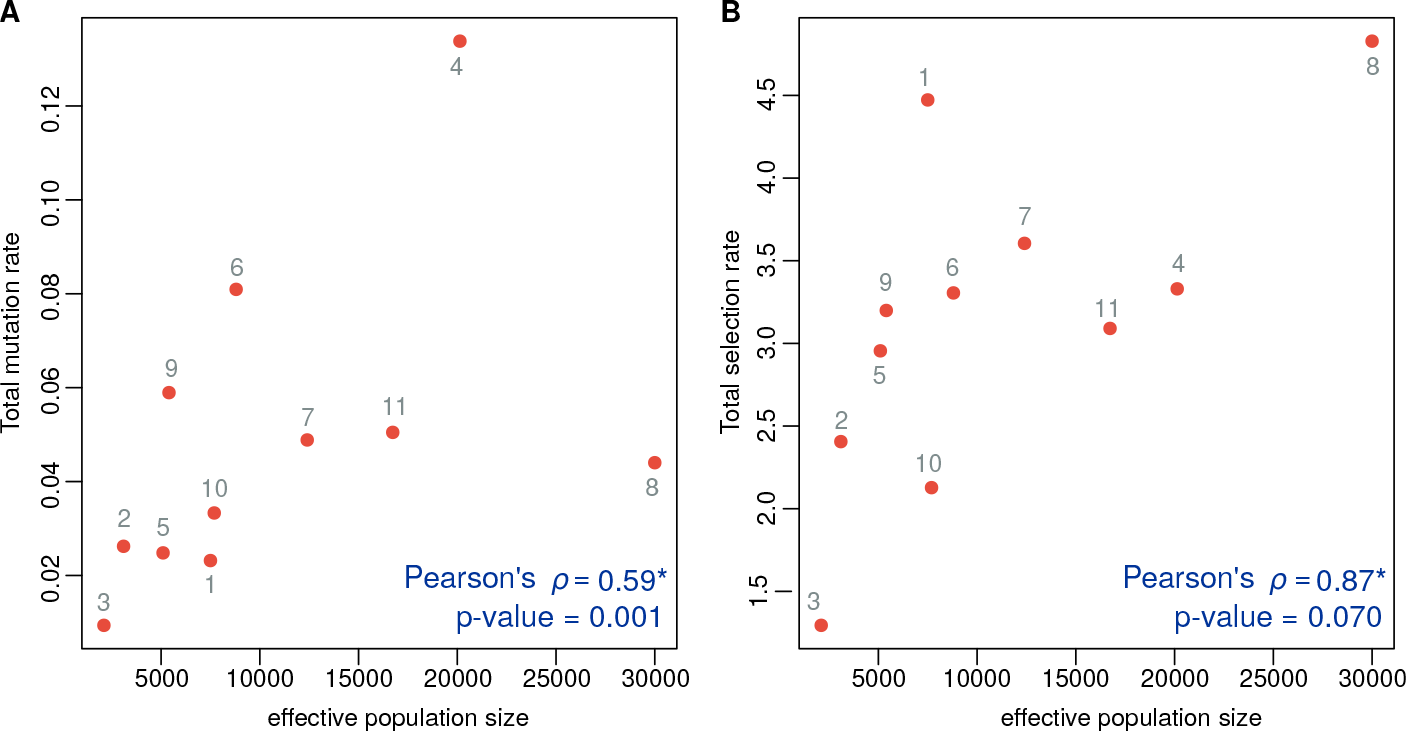
Correlating *N*_*e*_ with the total (A) rate of mutation and (B) selection in great apes. Great ape populations are numbered: 1. African humans, 2. Non-African humans, 3. Eastern gorillas, 4. Western gorillas, 5. Western chimpanzees, 6. Nigeria-Cameroon chimpanzees, 7. Eastern chimpanzees, 8. Central chimpanzees, 9. Bonobos, 10. Bornean orangutans and 11. Sumatran orangutans. Estimates of *N*_*e*_ were taken from Prado-Martinez et al. (2013) and Tenesa et al. (2007). * correlation coefficients calculated using independent contrasts and correcting for the effect of the great apes phylogeny (as predicted in Prado-Martinez et al. (2013))

This result shows that *N*_*e*_ plays an important role in determining the intensity of selection. In particular, it becomes clear that the different patterns of GC-bias found among great apes are, in part, due to different demographies. For example, Central chimpanzees have the highest GC-bias among the studied great apes, and they are indeed the population that was estimated with the largest *N*_*e*_ (30 000, Prado-Martinez et al. (2013)). Eastern gorillas showed the opposite pattern: this population had no evidence of GC-bias (with very homogeneous selection coefficients) and congruently Prado-Martinez et al. (2013) estimated its *N*_*e*_ as only 2000, the lowest of the studied populations.

**Table 2:**
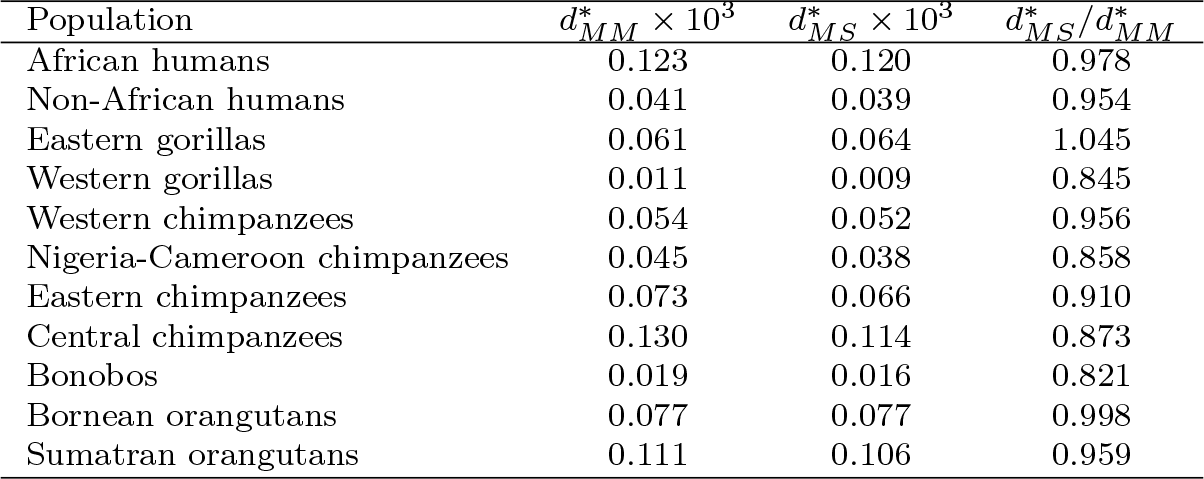
Expected number of substitutions per unit of time. The expected number of substitutions for the 4-variate Moran model with boundary mutations 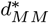 and allelic selection 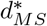 were calculated based on the posterior distributions of the model parameters and equation (6). The relative difference between the average number of events between the two models 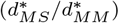 was used to assess how dissimilar these distances are.

| Population | $d_{MM}^* \times 10^3$ | $d_{MS}^* \times 10^3$ | $d_{MS}^*/d_{MM}^*$ |
| --- | --- | --- | --- |
| African humans | 0.123 | 0.120 | 0.978 |
| Non-African humans | 0.041 | 0.039 | 0.954 |
| Eastern gorillas | 0.061 | 0.064 | 1.045 |
| Western gorillas | 0.011 | 0.009 | 0.845 |
| Western chimpanzees | 0.054 | 0.052 | 0.956 |
| Nigeria-Cameroon chimpanzees | 0.045 | 0.038 | 0.858 |
| Eastern chimpanzees | 0.073 | 0.066 | 0.910 |
| Central chimpanzees | 0.130 | 0.114 | 0.873 |
| Bonobos | 0.019 | 0.016 | 0.821 |
| Bornean orangutans | 0.077 | 0.077 | 0.998 |
| Sumatran orangutans | 0.111 | 0.106 | 0.959 |

### 3.4 Comparing the expected number of substitutions in great apes

We calculated the expected number of substitutions under *MM* and *MS* to evaluate the impact of allelic selection (in particular, GC-bias) in the evolutionary process. With equation (6), we calculated 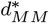 and 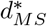 using the posterior estimates of the respective model parameters. We observe that for most of the great ape populations, the expected number of substitutions is lower when allelic selection is accounted (Table 2); Eastern gorillas are an exception, and the opposite pattern was observed. We also calculated the ratio between the expected number of substitutions in both models (i.e. 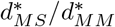) and we obtained minor (99.8% in Bornean orangutans) to major (82.1% in bonobos) deviations; the average difference is −7.3% (Table 2). These results suggest that not accounting for GC-bias may distort the reconstructed evolutionary process by overestimating the expected number of substitutions.

We complement this result by comparing the posterior distribution of the mutations rates in *MM* and *MS*. Because we wanted to identify the mutational types that may be differently estimated between these models, we calculated the relative difference between the mutation rate from allele *a*_*i*_ to allele *a*_*j*_ usiung the following ratio: 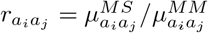. If 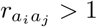 for a certain mutation rate *a*_*i*_*a*_*j*_, then this mutation rate is being underestimated in *MM* when compared to *MS* (and *vice versa* if 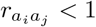); if 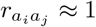 the mutation rates are equally estimated in both models.

We observed a systematic bias among great apes. While weak-to-weak and strong-to-strong mutation rates are generally non-deferentially estimated in both models (most of their *r* overlap 1, Figure 4) the strong-to-weak and weak-to-strong mutation rates are generally biased in *MM*. In particular, we obtained that weak-to-strong mutation rates are augmented, while mutations rates from strong-to-weak alleles are deprecated (Figure 4), which suggests that not accounting for GC-bias may bias the estimation of mutation rates. Eastern gorillas behave differently by not showing significant differences between the estimated mutations rates (all 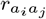 overlap 1, Figure 4).

**Figure 4:**
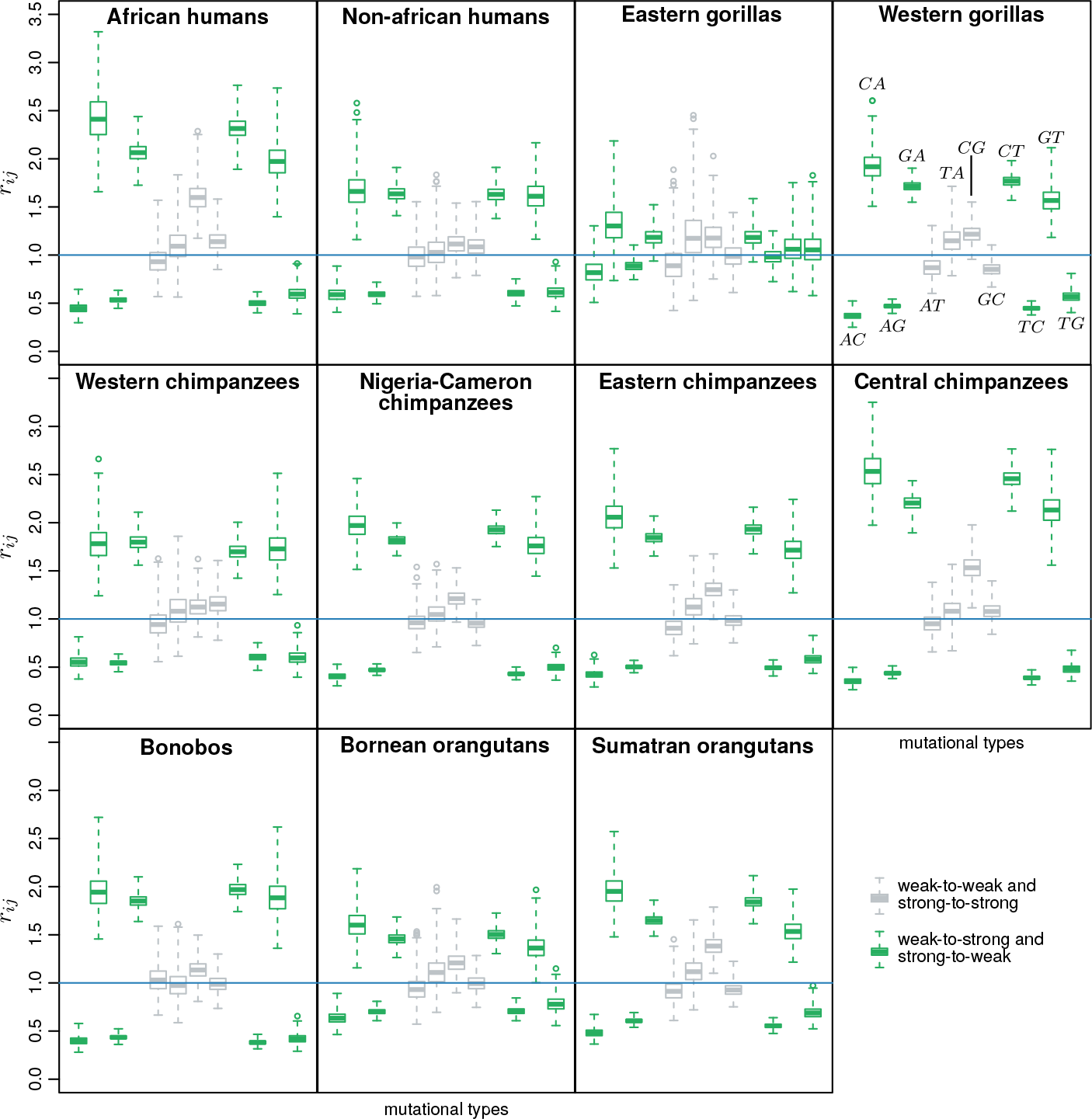
Relative difference in the mutation rates estimated under the neutral and non-neutral Moran model. 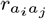 represents the ratio between the mutation from allele *a*_*i*_ to allele *a*_*j*_ in the model with allelic selection and the model with boundary mutations: 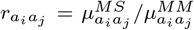. The 12 mutational types are indicated in the western gorillas plot: all of the plots follow this arrangement.

## 4 Discussion

In this work, we built on the multivariate Moran model with boundary mutations and allelic selection to explain the population processes shaping the observed distribution of alleles. We obtained new formulae to characterize this model. In particular, we derived the stationary distribution and the rate of the process. In addition, we built a Bayesian framework to estimate population parameters (mutation rates and selection coefficients) from population data. This work accomplishes tasks set by Schrempf and Hobolth (2017) who observed derivations from neutrality without having a model in place to enlighten the causes.

### 4.1 Variable patterns of gBGC among great apes

A genome-wide application to the great apes provides important insight into the strength and magnitude of GC-bias patterns and also the impact of gBGC in the evolutionary process. To our knowledge, this is the first work giving a population perspective of the patterns of GC-bias in non-human populations.

Here, we focus on GC-bias because it is a genome-wide effect. Mathematically speaking, it is difficult to disentangle gBGC from directional selection: they may have different biological explanations, but represent the exact same process modeling-wise (i.e. one allele is preferred over the others). Therefore, existing signatures of directional selection are most likely canceling out, when several site-histories (around 2.8 million sites in our case) are summarized to perform inferences.

In agreement with previous studies in mammals and humans (Spencer et al., 2006; Lartillot, 2013; Capra et al., 2013; Lachance and Tishkoff, 2014; Glémin et al., 2015), we found that gBGC is weak on average. Indeed, among great apes, the effect of GC-bias ranges between 1.49 *±* 0.53 (valued obtained by averaging *σ*_*C*_ and *σ*_*G*_), consistent with the nearly-neutral scenario (Ohta and Gillespie, 1996; Vogl and Bergman, 2015). These estimates are in congruence with other estimates of the scaled conversion coefficient in coding regions: Lynch (2010) estimated 4*N*_*e*_*s* as 0.82 in humans and Lartillot (2013) adopted a phylogenetic approach that predicted scaled conversion coefficients lower than 1 in all apes. The latter works employed the Wright-Fisher model; because we employed the Moran model, which has a rate of genetic drift twice as fast as the Wright-Fisher model, we expect to estimate twice as high selection coefficients.

We did not find a quantitative agreement between our estimates of the gBGC rate coefficient and those of the method of Glémin et al. (2015). In addition, we found that our model attributes to mutation what Glémin et al. (2015) attributes to gBGC. This might be a consequence the use of monomorphic sites by our method. Indeed, our estimates of gBGC correlate positively with the percentage of fixed sizes, but not those of Glémin et al. (2015). In general, the gBGC rate coefficient should promote more fixations by boosting the purging of polymorphic sites (at least for low mutation rates, as those observed in humans). On the other hand, Glémin et al. (2015) also considered a varying GC-content, which may explain why their estimates of gBGC are not correlating with the percentage of fixed sites. We have preliminary evidence showing that monomorphic sites can significantly impact estimates of population parameters. Nevertheless, a more comprehensive model accounting for both fixed states and variable GC-content would be necessary to disentangle their relative contribution to explaining the allele counts.

The patterns of GC-bias we have found in great apes are in concordance with the well-known process of gBGC. As expected, we observed that the larger the recombination rate or the lower the chromosome length, the higher the GC-effect. Evidently, recombination promotes gBGC; however, a negative association between gBGC and chromosome size is expected (in most organisms, small chromosomes undergo more recombination per unit of physical distance than large chromosomes (Kaback et al., 1989)). We have performed these analyses in non-African Humans, for which this data is available; however, we are confident that the patterns of GC-bias found in great apes are due to gBGC.

It has been hypothesized that GC-bias is a compensation mechanism for the mutational bias that exists in favor of the weak alleles, A and T (Galtier et al., 2009; Duret and Galtier, 2009; Philippe et al., 2011). Congruently with this expectations, we observed that mutation rates from strong to weak alleles are higher but rather similar between alleles of the same type. Interestingly, this symmetric manner by which mutations and selection are acting in great apes leads, as we have demonstrated, the number of substitutions to decrease in average. This suggests that the AT/GC toggling may increase the population variability by promoting more polymorphic sites, however, further studies would be necessary to clarify this prediction.

### 4.2 Intensity of gBGC and demography in great apes

Glémin et al. (2015) hypothesized that differences in GC-bias intensity among human populations were due to effects of demography. We also advance that demography regulates the intensity of gBGC in great apes. We obtained a positive correlation between the total rate of selection and *N*_*e*_ in great apes. An important conclusion of our study is that the patterns of gBGC can rapidly change due to demography, even among closely related populations. In fact, most of the studied populations are known to have diverged less than 0.5 million years ago (Prado-Martinez et al., 2013).

Here, we showed that GC-bias determines the genome-wide base composition of genomes in a factor proportional to (1 + *σ*_*C/G*_)^*N* − 1^ (or 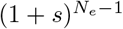 in the true dynamic). Therefore, by either changing *N*_*e*_ or *s*, we are able to change the AT/GC composition of genomes. Because we were able to correlate *N*_*e*_ with the intensity of allelic selection (Pearson’s *ρ* = 0.87), we are convinced that demography has a major role determining the base composition of great apes genomes. Studies using life history traits (i.e. body size) in mammals (Romiguier et al., 2010) and ancestral reconstructions of the effective population size in birds (Weber et al., 2014) also advocated for correlations between *N*_*e*_ and GC-content (although not so strong as the one found here; *ρ* around 0.30 − 0.55 in Weber et al. (2014))

In contrast, Galtier et al. (2018) have not found this correlation in a data set covering 31 species of distinct metazoa phyla (including vertebrates, insects, molluscs, crustaceans, echinoderms, tunicates, annelids, nematodes, nemertians and cnidarians). This is most likely happening because aspects of the recombination landscape may also affect the intensity of gBGC (Duret and Galtier, 2009; Lesecque et al., 2014; Galtier et al., 2018): genome-wide recombination rate, length of gene conversion tracts and repair biases. As the recombination landscape significantly varies across species, but not so much across related populations (e.g. the caryotype is very conserved among great apes, with humans having 46 diploid chromosomes whereas other great apes having 48), we expected stronger correlations between the intensity of gBGC and demography.

Knowing to what extent variations in *N*_*e*_ or *s* determine the base composition of genomes will require further studies. In particular, determining *s* experimentally in different populations/species would help to assess the real impact of gBGC. If we could assume that *s* vary slightly among closely related populations/species, then we might attribute different intensities of GC-bias almost solely to demographic effects, which simplifies the task of accommodating gBGC in population models.

### 4.3 gBGC calls for caution in molecular and phylogenetic analyses

The effects of gBGC in the molecular analysis have been extensively described in the literature (reviewed in Romiguier and Roux (2017)), we complement these results by showing how GC-bias affects the base composition of genomes, and how the mutation rates and genetic distances may be biased if GC-bias is not properly accounted. In particular, we observed that mutations rates from weak-to-strong and strong-to-weak alleles are systematically over and underestimated, respectively.

The idea that gBGC may distort the reconstructed evolutionary process comes mainly from phylogenetic studies. For example, it is hypothesized that gBGC may promote substitution saturation (Romiguier and Roux, 2017). We have shown that the number of substitutions may be significantly overestimated if we do not account for GC-bias, meaning that gBGC may indeed promote branch saturation. Based on this and other gBGC-related complications (e.g. GC-bias promotes incomplete lineage sorting (Hobolth et al., 2011)), some authors advocate that only GC-poor markers should be used for phylogenetic analysis (McCormack et al., 2012; Romiguier et al., 2013). Contradicting this approach, our results show that we may gain more inferential power if GC-bias is accounted for when estimating evolutionary distances.

Here, we have not performed phylogenetic inference, but previous applications of the Moran model to phylogenetic problems (i.e. PoMo) (De Maio et al., 2015; Schrempf et al., 2016) show that it can be done. Therefore, a necessary future work would be testing the effect of allelic selection (or, more specifically, GC-bias) in phylogeny reconstruction; in particular, it would be of major interest determining how much of its signal can be accounted for increasing the accuracy of tree estimation.

Recently, a nucleotide substitution process that accounts for gBGC was proposed by Lartillot (2013). In this model, the scaled conversion coefficient is used to correct the substitution rates in a similar fashion as we have done to calculate the expected number of substitutions for the Moran distance (i.e. assessing the relative fixation probabilities under GC-bias, File S3). Therefore, these models should perform similarly, with the exception that PoMo should be able to disentangle the contribution of selection and mutation to the observed diversity, as it additionally accounts for polymorphic sites.

## 5 Conclusion

Despite the widespread evidence of gBGC in several taxa, several questions remain open regarding the role of gBGC determining the base composition of genomes. In this work, we provided a mechanistic model and theoretical results that allowed quantifying patterns of gBGC in non-human closely related populations that have given a new layer of understanding on the tempo and mode of gBGC evolution in vertebrate genomes.

In addition, our multivariate Moran model with allelic selection adds a significant contribution to the endeavor of estimating population parameters from multi-individual population-scale data. Importantly, our analysis showed that gBGC may significantly distort estimates of population parameters and genetic distances, stressing that gBGC-aware models should be used when employing molecular phylogenetics and population genetics analyses. We stress that although our application to great apes show evidence of GC-bias, our framework can be more generally employed to estimate patterns of nucleotide usage and associated mechanisms of evolution.

## Acknowledgements

This work has been funded by the Vienna Science and Technology Fund (WWTF) through project MA16-061. GS received funding from the European Research Council under the European Unions Horizon 2020 research and innovation program under grant agreement no. 714774. We thank Dominik Schrempf, Claus Vogl and Sylvain Glémin for helpful discussions, and the three anonymous reviewers for comments improving the manuscript.

## A Appendix: Virtual population size

Consider two populations *A* and *A*′ with different population size *N* and *M*, respectively. We want to mimic the dynamics of population *A*, relying on the population parameters of a population *A*′ of different size (larger or smaller). Both population have the same number of monomorphic states (equaling the number of alleles *K*) and so we assume them equally frequent in both populations. The number of polymorphic states differs: there are *K*(*N* − 1) polymorphic states in population *A*, while *A*′ has *K*(*M* − 1). Because we cannot make polymorphic states equivalent, we assume that the sum of polymorphic states for each pairwise comparison of the *K* alleles should be equal in both populations. These conditions can be written in the following system of equations

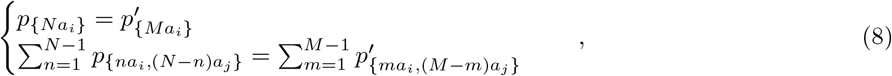

which will have *K* + *K*(*K* − 1/2) conditions in total.

As we have derived an estimator of the site frequency spectrum, we can write this conditions for the multivariate Moran model with boundary mutations and selection as

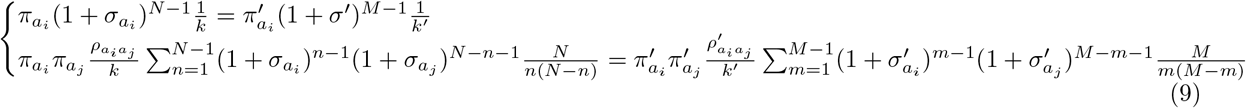

This system has *K* + *K*(*K* − 1)/2 conditions and 2*K* − 2 + *K*(*K* − 1)/2 parameters and therefore cannot be solved. However, we know that the entries of *π* are constrained in [0, 1] and should sum up to 1 in both populations, therefore we make the additional assumption that 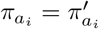. In addition, and by definition, the reference allele 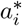 is considered to evolve neutrally in both systems, which permits to conclude that the normalization constants *k* and *k*′ are equal. Simplifying,

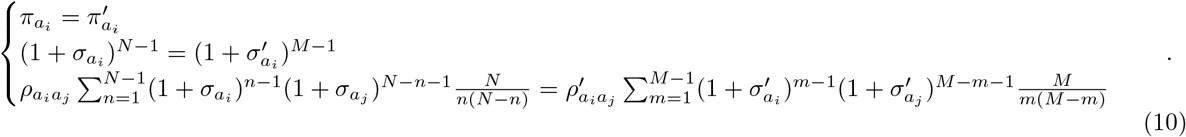

we obtain that the population parameters of population *A*′ can be expressed in terms of the parameters of population *A*

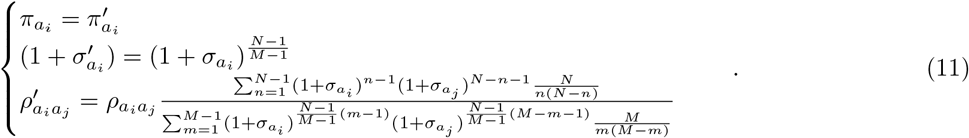

This expression looks tedious, but the neutral case 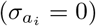 can be very intuitive. In this scenario, mutation rates of populations *A* and *A*′ change by a factor that is simply the ratio of two harmonic numbers, each of which determined by the population size of the respective population. Intuitively, if *N* > *M* then 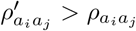, meaning that in order to compensate the smaller number of individuals *M* (i.e. stronger effect of genetic drift), mutation rates are augmented in population *A*′. Figure 5 depicts the effect of the effective population size on the mutation rates and selection coefficients.

**Figure 5:**
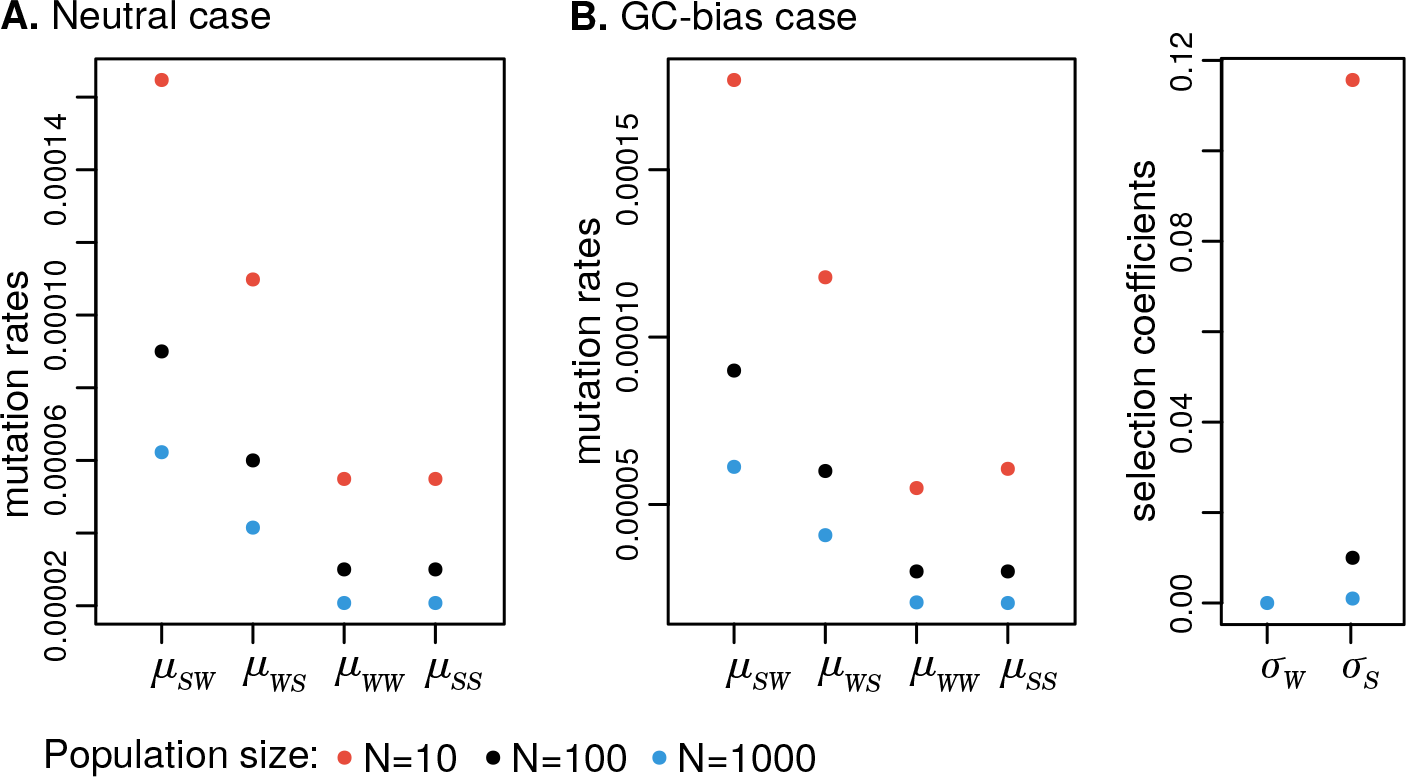
Population parameters transformation for different population sizes. We considered the simple case of two alleles: W stands for the weak alleles *A* and *T*, and S stands for the strong alleles *C* and *G*. Model parameters were set to represent a **A.** neutral case (black dots: *μ*_*SW*_ = 9 × 10^−5^, *μ*_*WS*_ = 6 × 10^−5^ and *μ*_*WW*_ = *μ*_*SS*_ = 3 × 10^−5^) and a **B.** GC-bias case (black dots: mutations rates equal to the neutral scenario and *σ*_*W*_ = 0, *σ* = 0.01).

## B Appendix: Proof of the stationary vector

Let *ψ* be a stationary vector of **Q** with 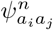 and *ψ*_*i*_ being the elements of the stationary vector corresponding to the states {*na*_*i*_, (*N* − *n*)*a*_*j*_} and {*Na*_*i*_}, respectively. In the multivariate Moran model with low mutation rates and selection, mutation is only occurring in the boundary states, permitting the monomorphic states to communicate with the polymorphic states, while drift and selection are both acting among the polymorphic states. The detailed balance conditions can be defined and lead to equations for the monomorphic and the polymorphic states. In the boundary states, an allele *a*_*i*_ is either fixed (*n* = *N*) or absent (*n* = 0, i.e. *a*_*j*_ is fixed), for which we may write

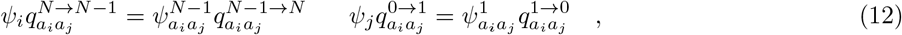

while between the polymorphic states, the general condition is valid

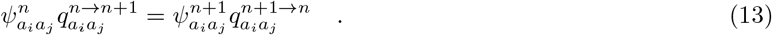

Condition (13) can be rewritten in the recursive form

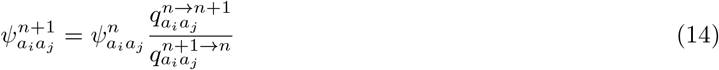

and then combined with equation (12)

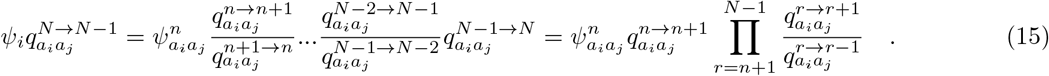

The product can be further simplified by recognizing that for *r* = *N* − 1, 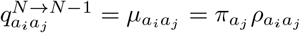, while for *r* < *N* − 1, the rates inside the product are just the combined rate of drift and selection (according to expression (2). We can now rewrite equation (14) in order to the 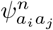 element of the stationary vector of **Q**

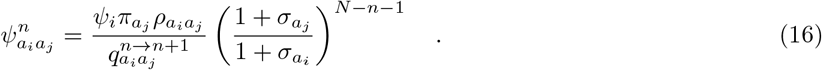

Because 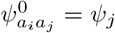 and 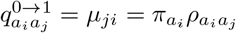, we obtain a possible solution for the monomorphic states of the stationary distribution by making *n* = 0 in equation (16)

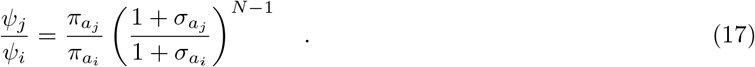

The stationary solution for the polymorphic states can be obtained from equation (16) by noting that 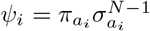 and 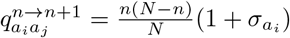

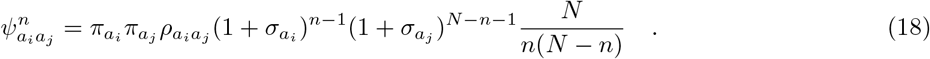

The stationary distribution obtained here can be related with the stationary vector of the neutral boundary multivariate Moran model. We observe that when ***σ*** = **0**, we obtain the solution computed by Schrempf et al. (2016) for the multivariate Moran model with drift only

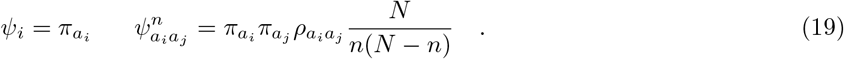

## C Appendix: Proof of the expected number of Moran events per unit of time

To assess the impact of allelic selection in branch length estimation (or the total rate of the process), we computed the expected number of events per unit of time for the multivariate Moran model with selection

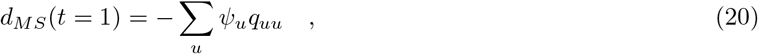

Where *ψ* is the stationary vector and *q*_*uu*_ the diagonal elements of ***Q***. Equation (20) can be solved by observing that a monomorphic state can only be escaped by mutation, while a polymorphic state can only be escaped by selection and drift

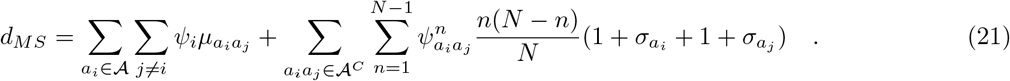

The stationary vector is known from equations (17) and (18)

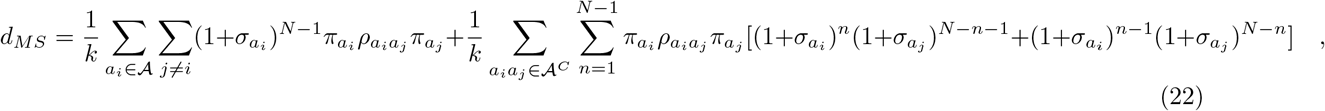

where *k* is the normalization constant defined in equation (4). Expression (22) can be further simplified by observing that the sum in *n* results in doubling every 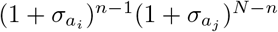 element. Therefore, the expected number of events can be simplified to

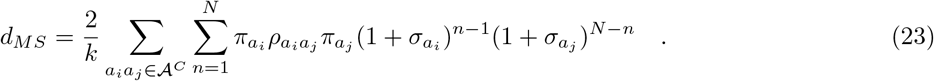

## D Appendix: Proof of the Moran distance in number substitutions

The Moran distance *d*_*MS*_ accounts for several events (mutation, drift and selection) and differs from the standard evolutionary distances because they are calculated in terms of the expected number of substitutions 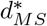.

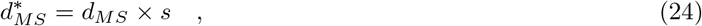

where *s* is the probability of a substitution. *s* can be calculated multiplying the probability *m* of an event being a mutation, by the probability *h* of that mutation getting fixed in the population

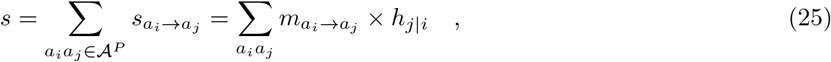

where 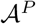 represents all the possible pair-wise permutations without repetition of *K* alleles.

## 1. Solving 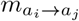

The probability of an event being a mutation is simply the ratio between the rate of mutation and the total rate (i.e the rate of mutation plus the rate of drift and selection). In stationarity, we know that the total rate *r*_*T*_ = *d*_*MS*_(1) is simply the expected number of events of the Moran model and follows equation (23). The rate of a *a*_*i*_ → *a*_*j*_ mutation is the rate of escaping the monomorphic state {*Na*_*i*_}, from which we can write

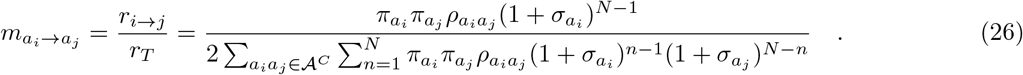

We can see that 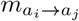 differs from 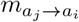 only due to the selection coefficient in the numerator.

## 2. Solving 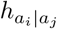

According to Kluth and Baake (2013), the fixation probability of an allele with fitness 1 + *σ* is for the Moran model

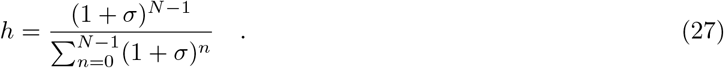

As we are using the multivariate Moran model, we have to extend the denominator of (27) to account for the different possible combinations of two selection coefficients. In particular, we may have

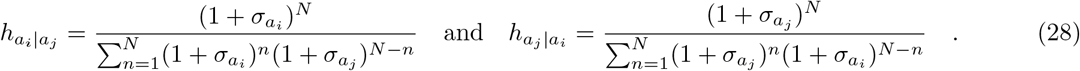

We further redefine the denominators in order to make them equal

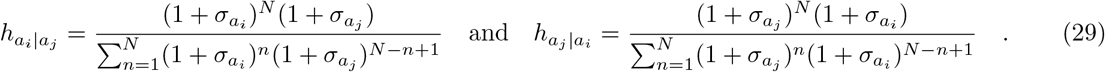

## 3. Solving *s*

The probability of a *a*_*i*_ → *a*_*j*_ substitution under the multivariate Moran model with boundary mutations and selection can be computed as

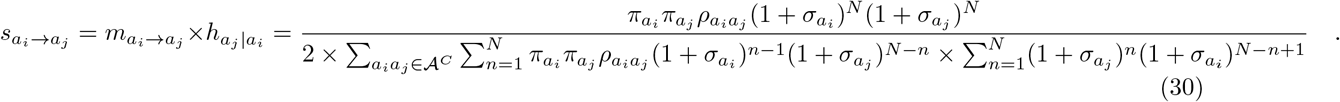

We see that 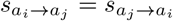, which is an expected consequence of stationarity. We can now generalize 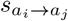 for all the substitution types by using equation (25)

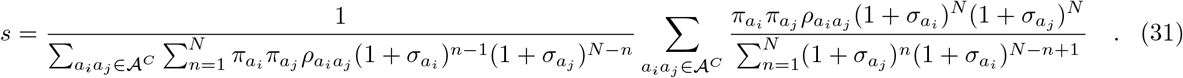

The relationship between the Moran distance in events and substitutions can be defined based on equation (24),

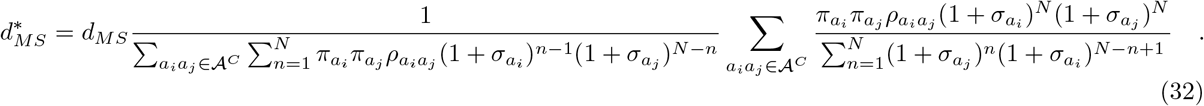

This quantity can be evaluated for neutral regimes: i.e. ***σ*** → (0, 0, 0, 0). We obtain the probability of a substitutions under the neutral Moran model and it matches the result computed by Schrempf et al. (2016):

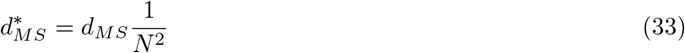

