## Supplementary figures and tables for "Quantifying GC-biased gene conversion in great ape genomes using polymorphism-aware models"

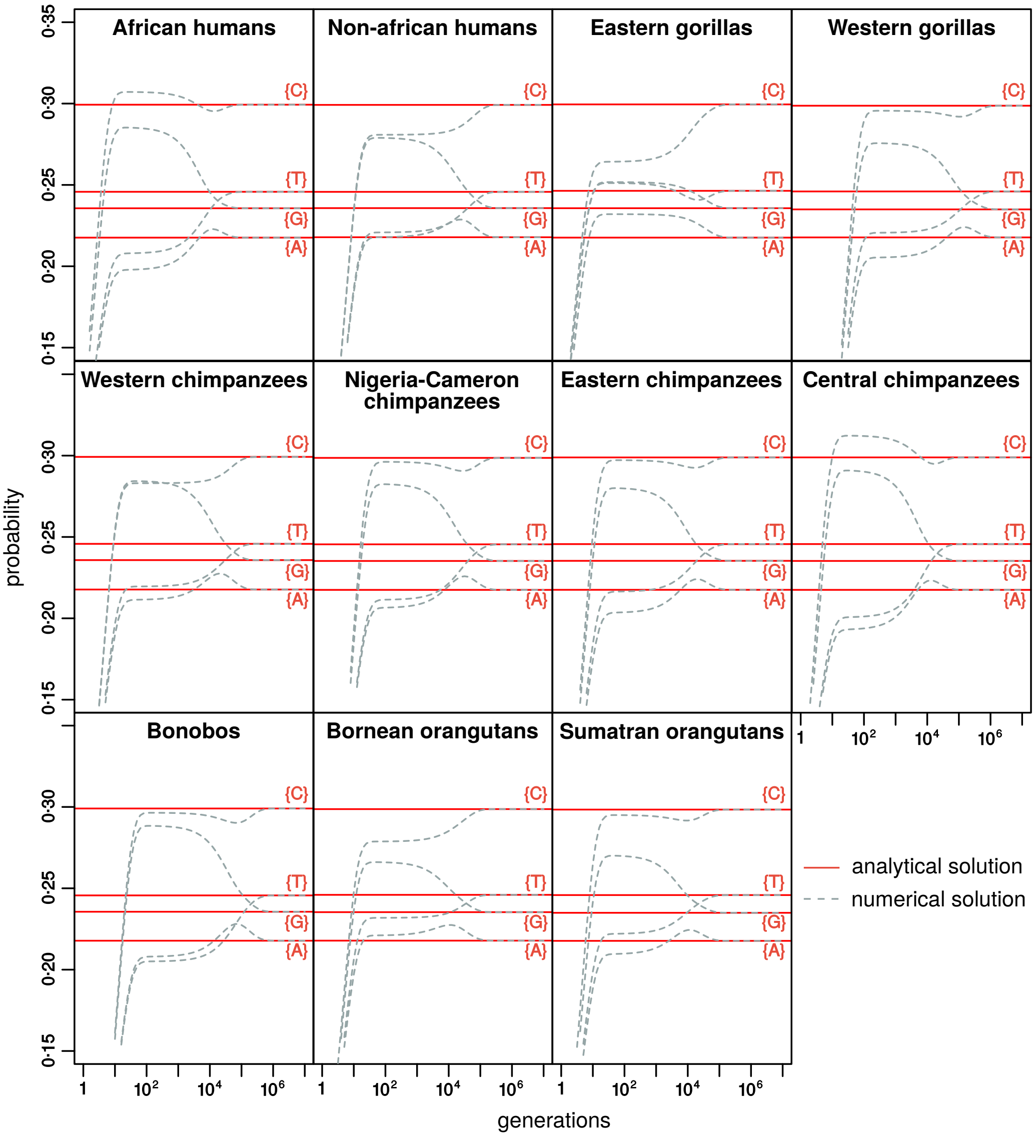


Figure S1: **Numerical validation of the stationarity vector**. Estimated vectors of $\boldsymbol{\pi}$, $\boldsymbol{\rho}$, $\boldsymbol{\sigma}$ from the great apes' data were used to calculate the rate matrix $\boldsymbol{Q}$ and the probabilities for the state space at several time points (time in generations). The initial probabilities were set uniformly as $\frac{1}{4+6(N-1)}$, i.e. the number of states. For sake of clarity only the monomorphic states $\{i\}$ are represented.

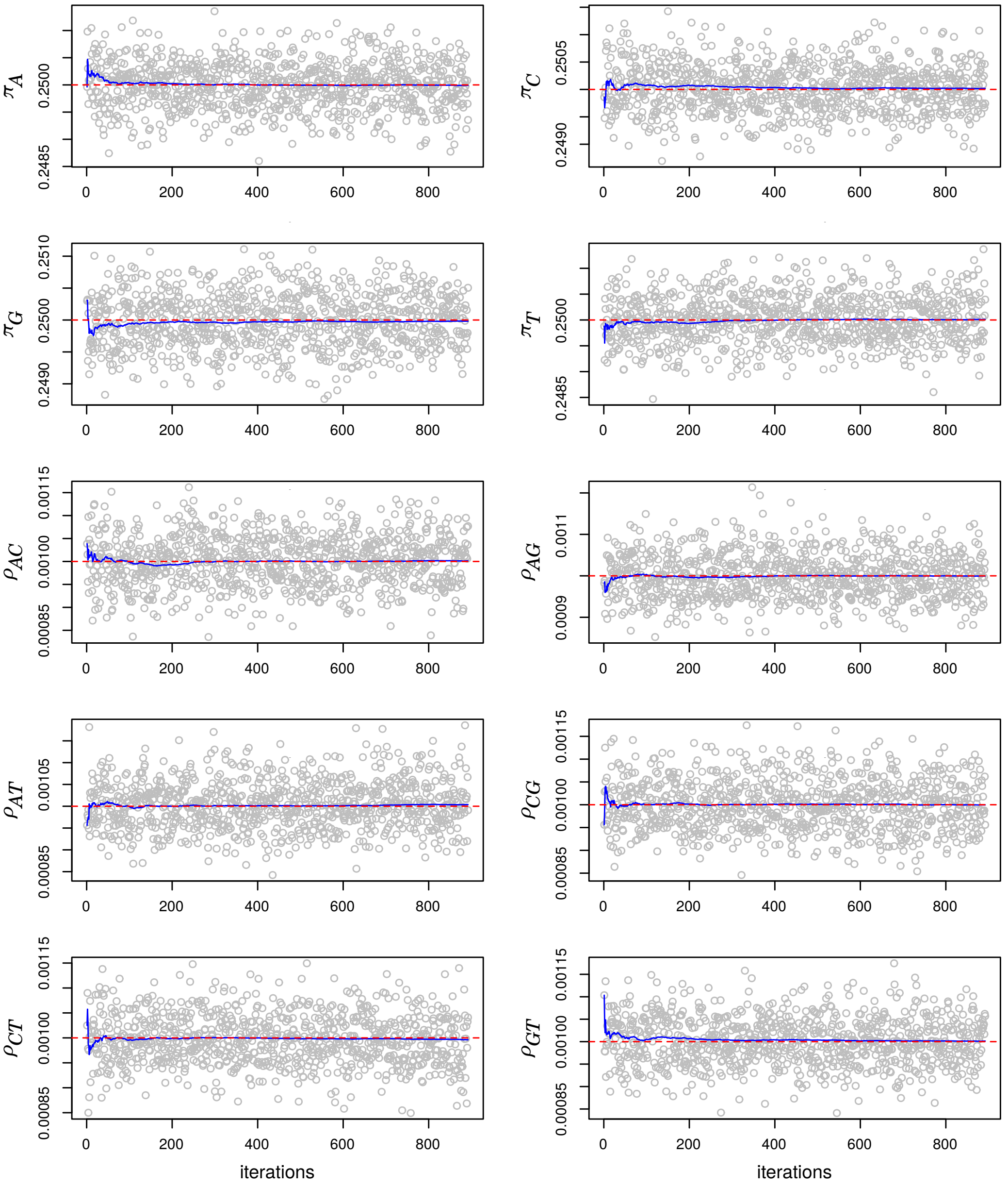


Figure S2: **Validation of the Bayesian algorithms.** Trace plot depicting the convergence of the MCMC runs (grey dots and blue lines) to the true parameter values (red lines). Simulation conditions: 1000000 sites, 10 individuals and a simple parameter vector for the Moran model with boundary mutations: $\boldsymbol{\pi} = (0.25, 0.25, 0.25, 0.25)$, $\boldsymbol{\rho} = (0.001, 0.001, 0.001, 0.001, 0.001, 0.001)$. The blue line represents the MCMC moving average whereas the red one represents the true values. The codes used to performed these simulations is available in GitHub: `https://github.com/pomo-dev/pomo_selection`.

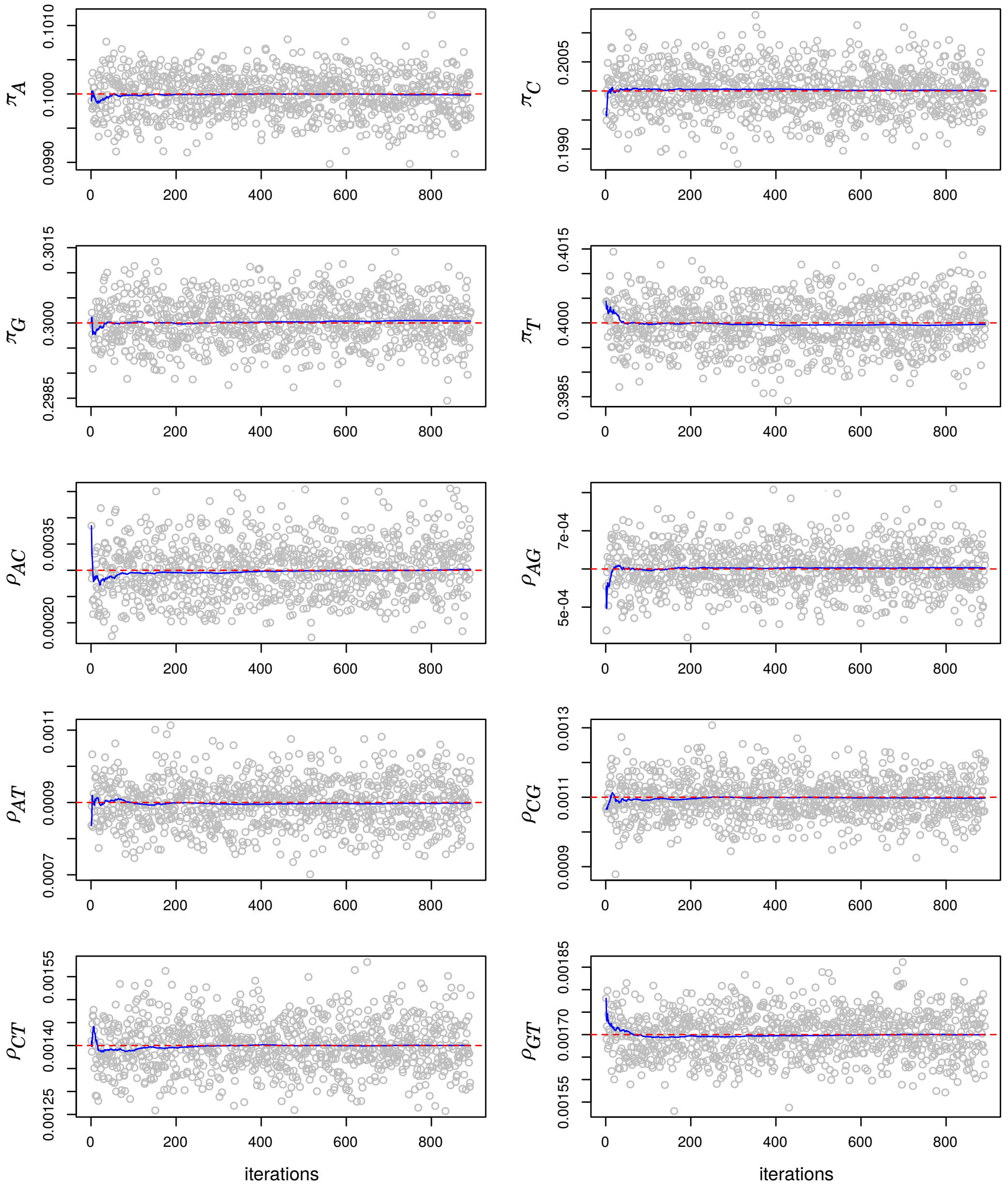


Figure S3: **Validation of the Bayesian algorithms.** Trace plot depicting the convergence of the MCMC runs (grey dots and blue lines) to the true parameter values (red lines). Simulation conditions: 1000000 sites, 10 individuals and a complex parameter vector for the Moran model with boundary mutations: $\boldsymbol{\pi} = (0.10, 0.20, 0.30, 0.40)$, $\boldsymbol{\rho} = (0.0003, 0.0006, 0.0009, 0.0011, 0.0014, 0.0017)$. The blue line represents the MCMC moving average whereas the red one represents the true values. The codes used to performed these simulations is available in GitHub: `https://github.com/pomo-dev/pomo_selection`.

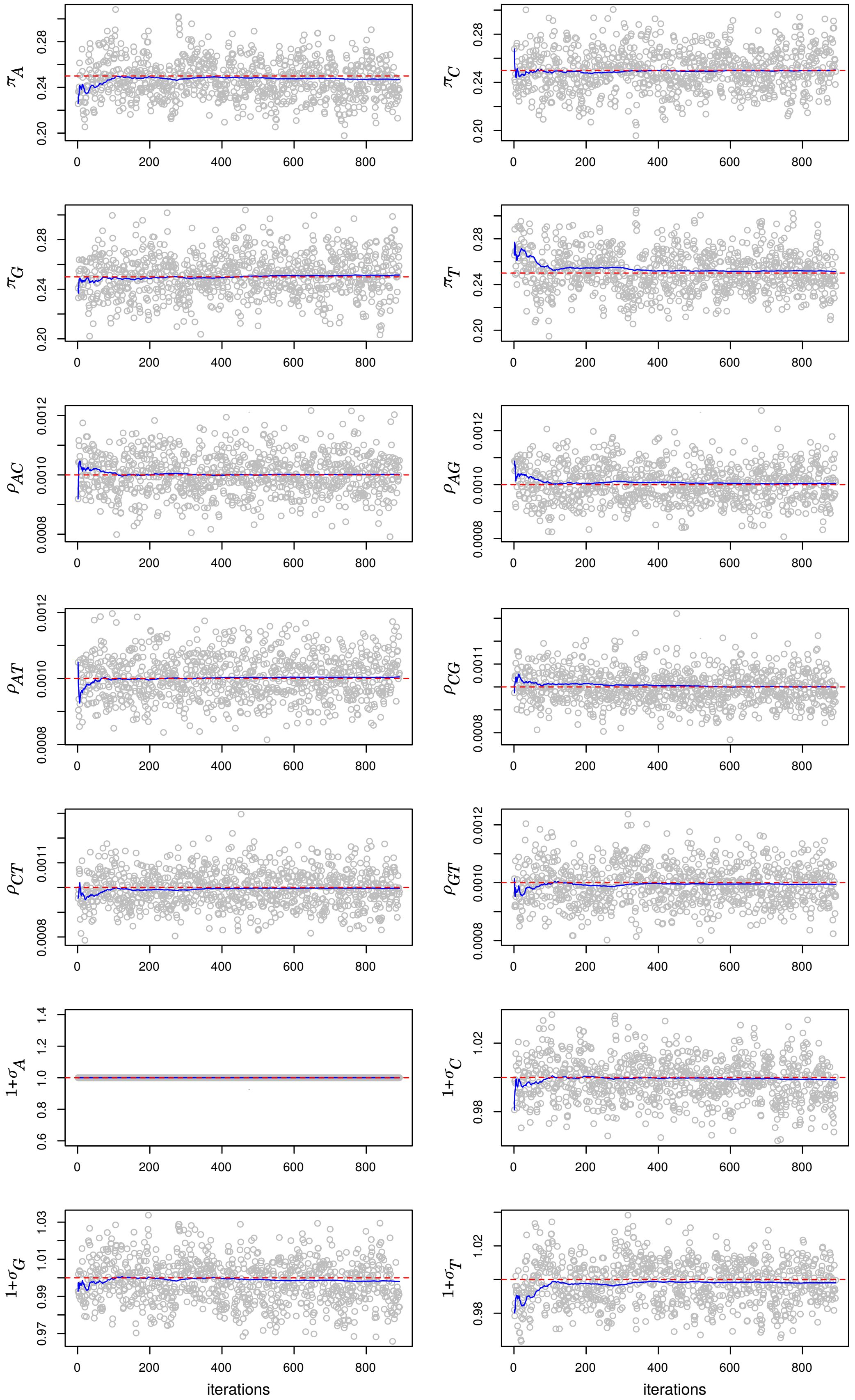


Figure S4: **Validation of the Bayesian algorithms.** Trace plot depicting the convergence of the MCMC runs (grey dots and blue lines) to the true parameter values (red lines). Simulation conditions: 1000000 sites, 10 individuals and a simple parameter vector for the Moran model with allelic selection: $\boldsymbol{\pi} = (0.25, 0.25, 0.25, 0.25)$, $\boldsymbol{\rho} = (0.001, 0.001, 0.001, 0.001, 0.001, 0.001)$, $\boldsymbol{\sigma} = (1.00, 1.00, 1.00, 1.00)$. The blue line represents the MCMC moving average whereas the red one represents the true values. The codes used to performed these simulations is available in GitHub: `https://github.com/pomo-dev/pomo_selection`.

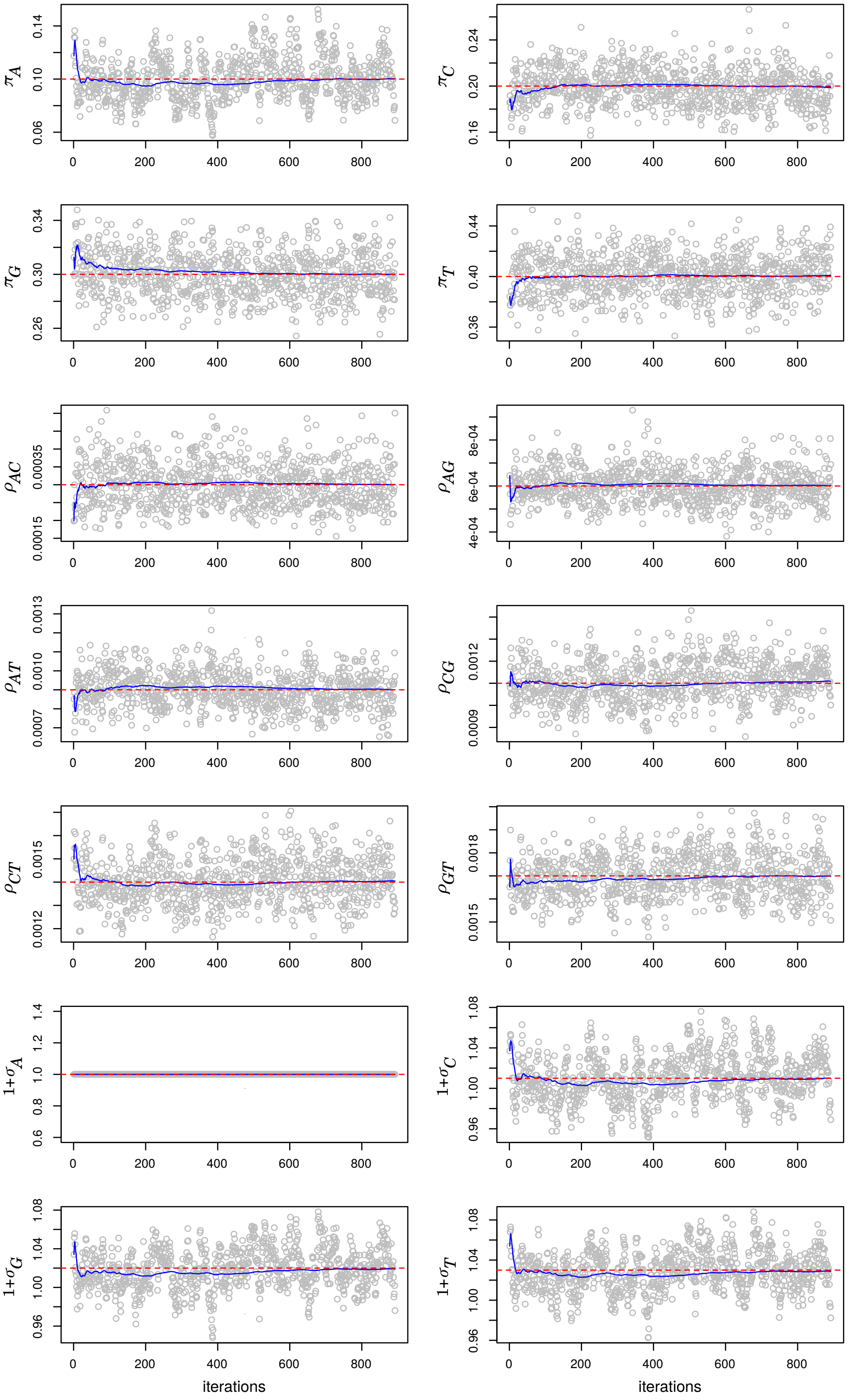


Figure S5: **Validation of the Bayesian algorithms.** Trace plot depicting the convergence of the MCMC runs (grey dots and blue lines) to the true parameter values (red lines). Simulation conditions: 1000000 sites, 10 individuals and a complex parameter vector for the Moran model with allelic selection: $\boldsymbol{\pi} = (0.10, 0.20, 0.30, 0.40)$, $\boldsymbol{\rho} = (0.0003, 0.0006, 0.0009, 0.0011, 0.0014, 0.0017)$, $\boldsymbol{\sigma} = (1.00, 1.01, 1.02, 1.03)$. The blue line represents the MCMC moving average whereas the red one represents the true values. The codes used to performed these simulations is available in GitHub: `https://github.com/pomo-dev/pomo_selection`.

Figure S6: **Prediction of the site-frequency spectrum in great ape populations.** The gray points represent the observed counts and the vertical lines the posterior predictive distribution of the stationary distribution under the 4-variate Moran model: boundary mutation model (blue) and allelic selection model (red).

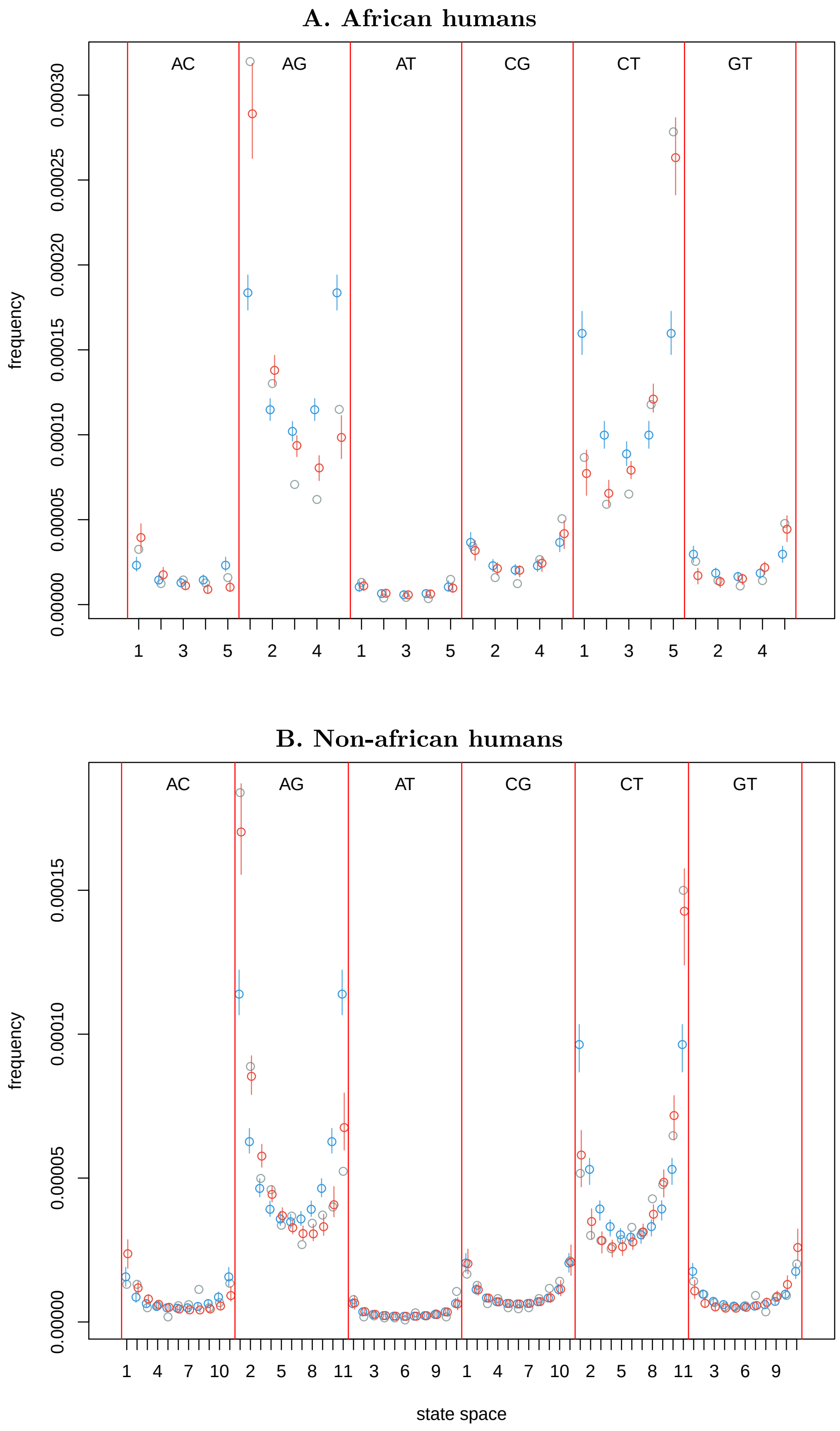

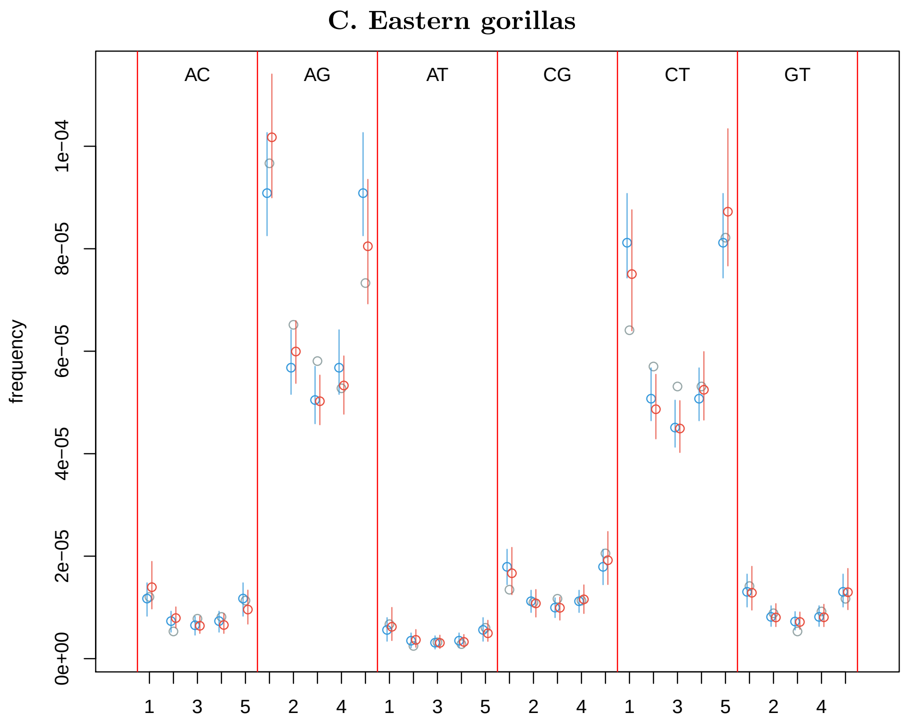
C. Eastern gorillas
AC
AG
AT
CG
CT
GT
frequency
0e+00
2e−05
4e−05
6e−05
8e−05
1e−04
1
3
5
2
4
1
3
5
2
4
1
3
5
2
4

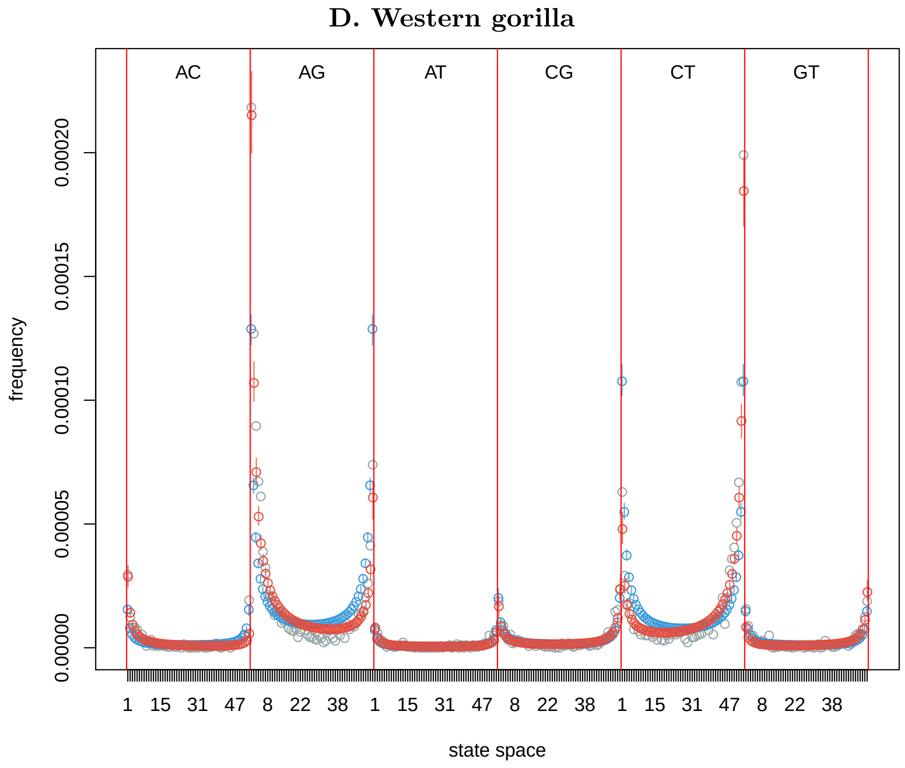
D. Western gorilla
AC
AG
AT
CG
CT
GT
frequency
0.00000
0.00005
0.00010
0.00015
0.00020
1
15
31
47
8
22
38
1
15
31
47
8
22
38
1
15
31
47
8
22
38
state space

**E. Western chimpanzees**

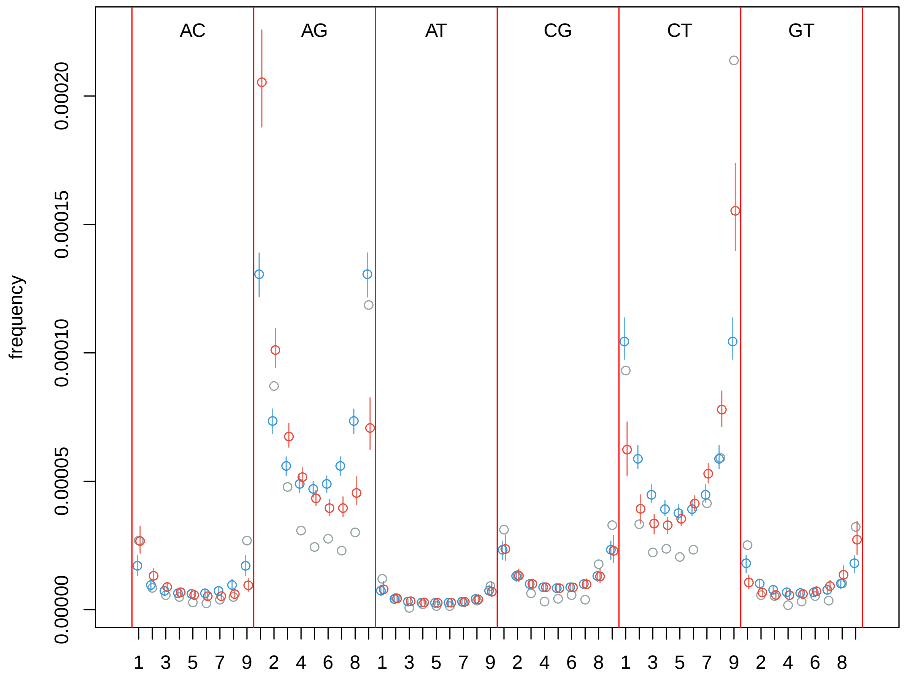


**F. Nigeria-Cameroon chimpanzees**

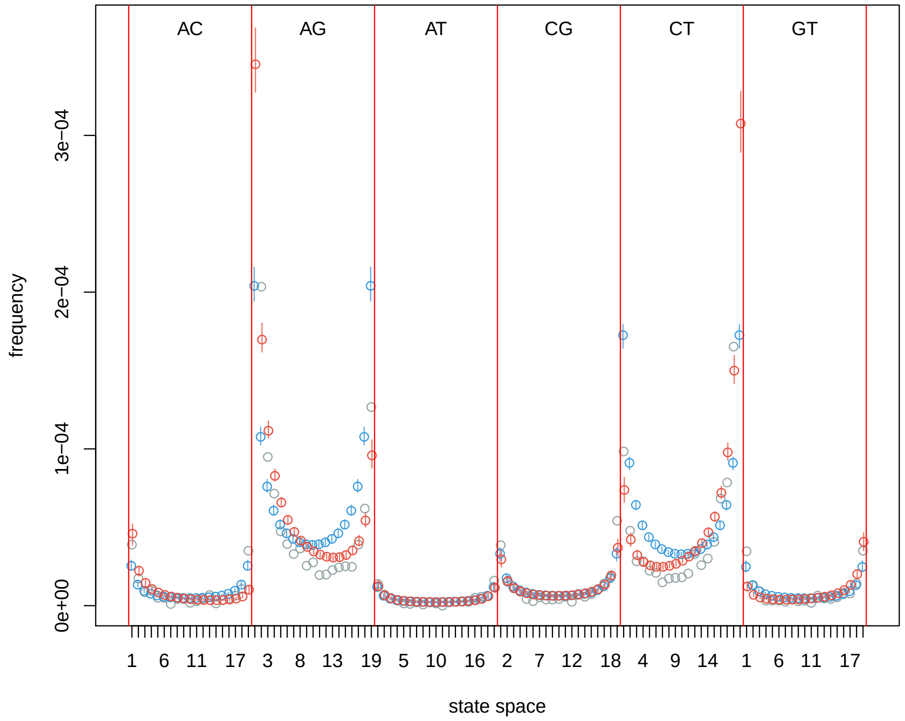

G. Eastern chimpanzees

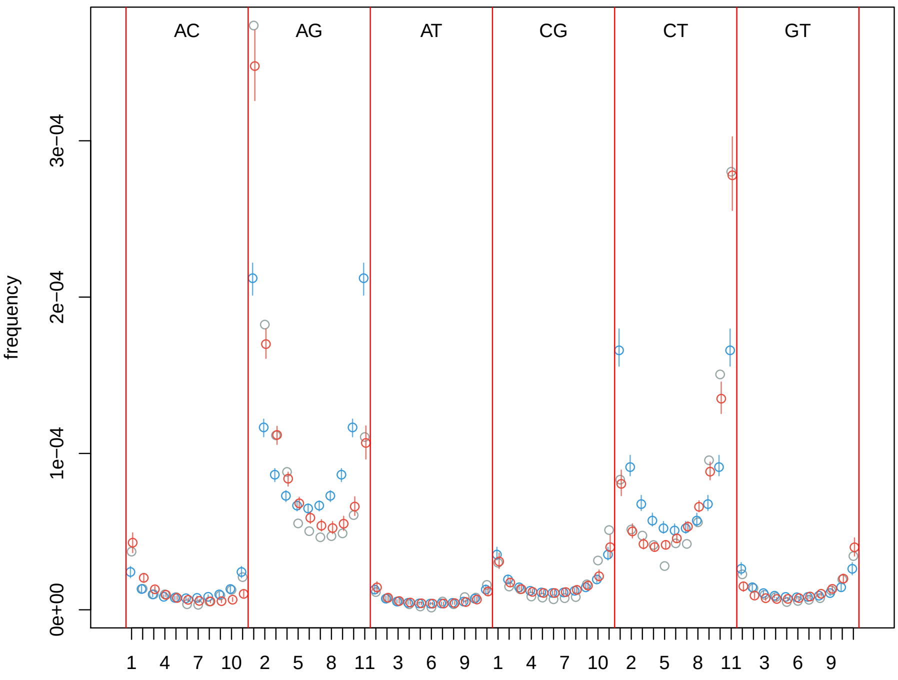


H. Central chimpanzees

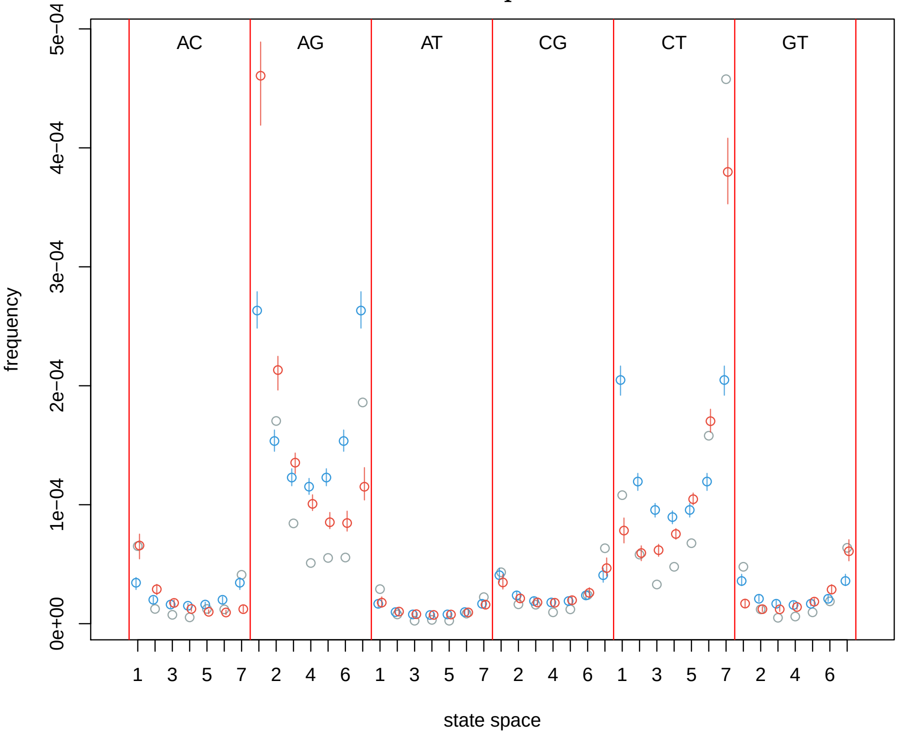

### I. Bonobos

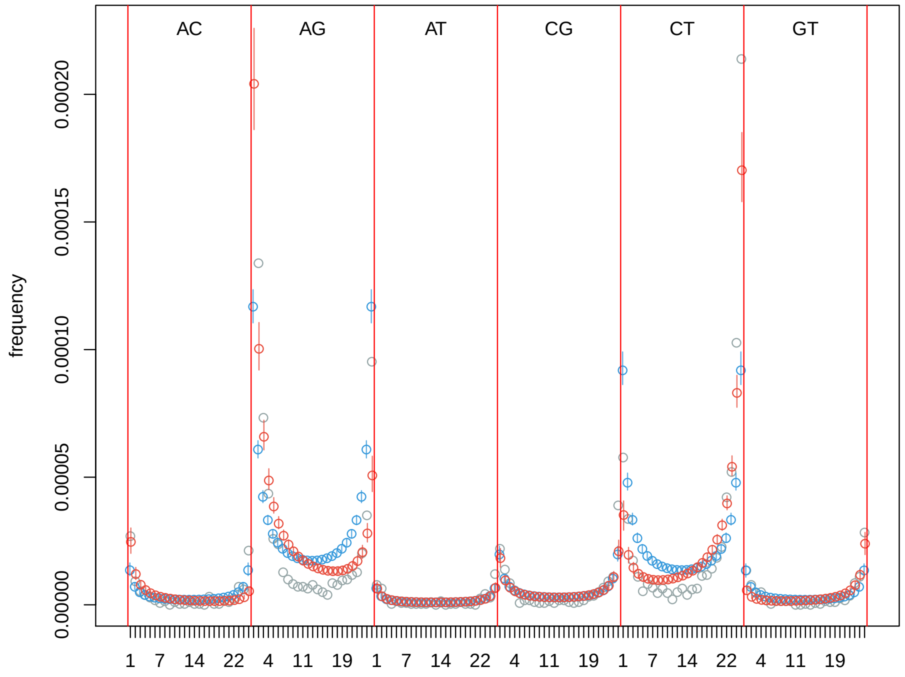


### J. Bornean orangutans

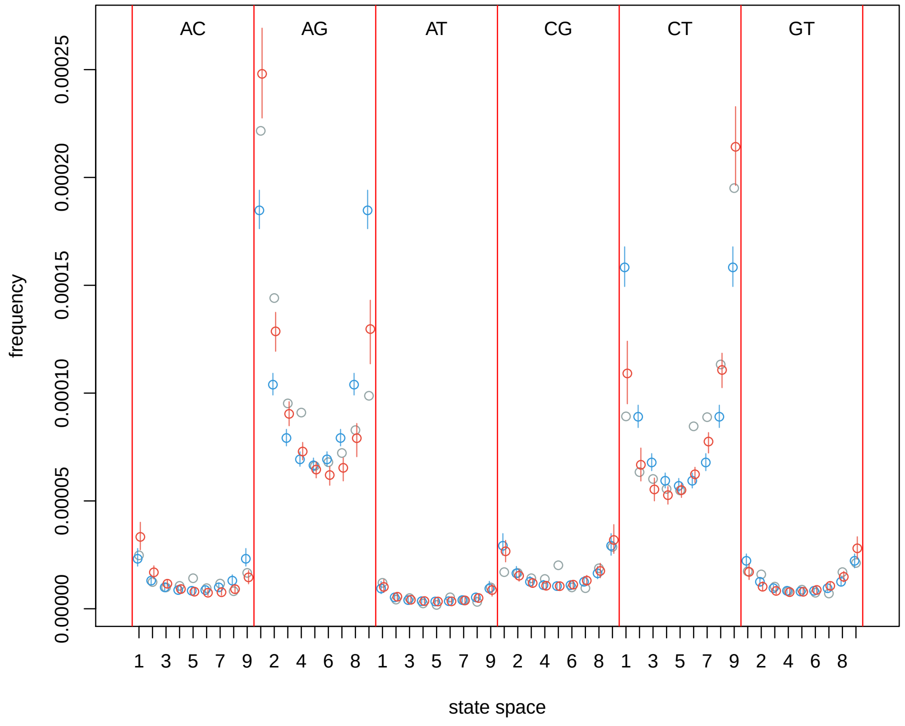

**K. Sumatran orangutans**

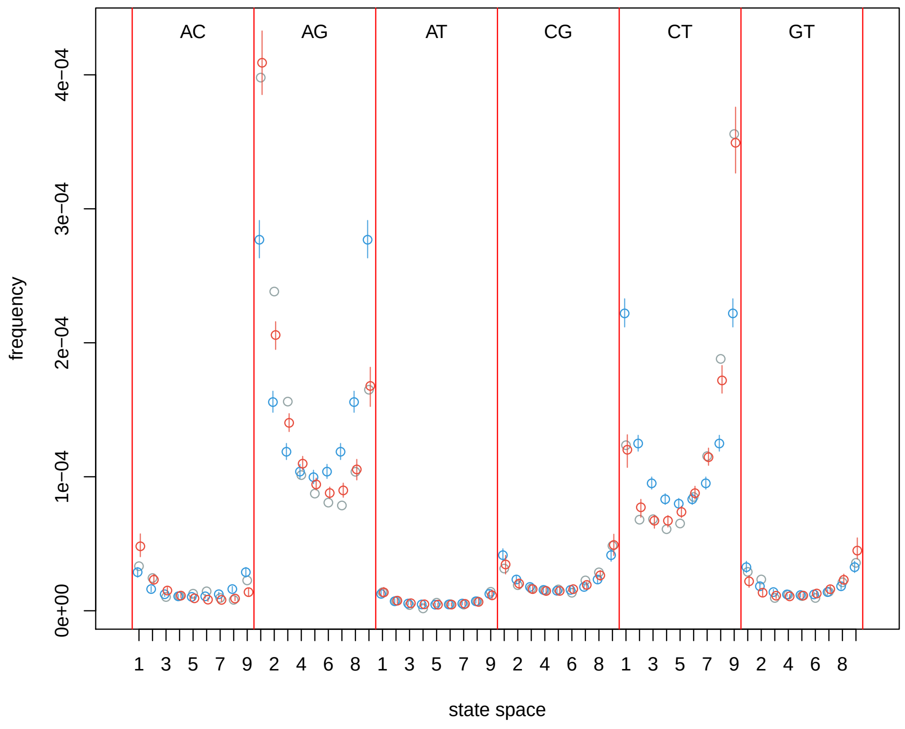

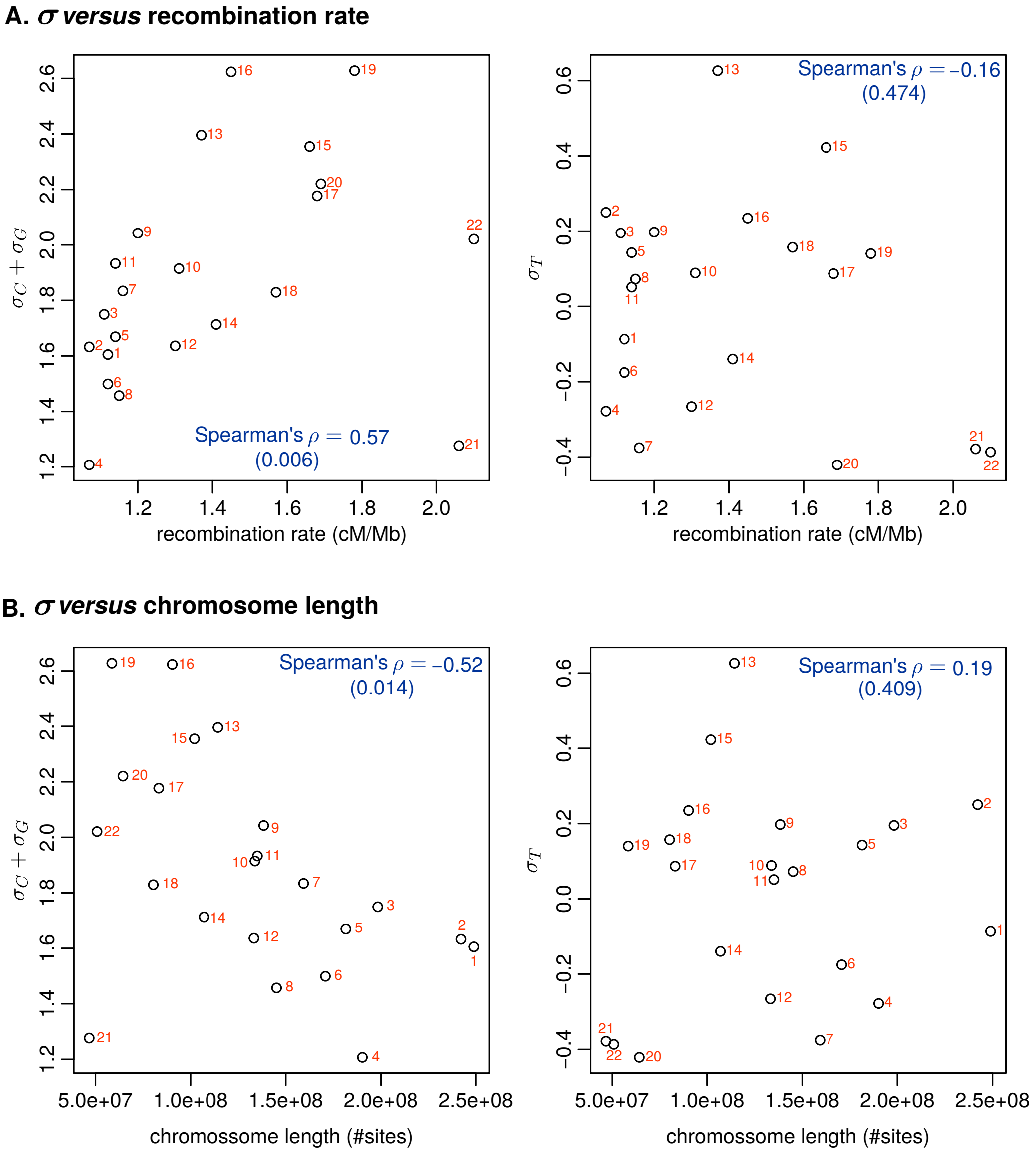


Figure S7: **GC-bias *vs.* recombination rate and chromosome length in non-African humans**. The scaled selection coefficients were estimated based on the posterior average. Recombination rates were estimated by comparing the genetic distance (cM) between markers to the physical (Mb) as described in (Jensen-Seaman (2004)) and based on the human Iceland pedigree map.

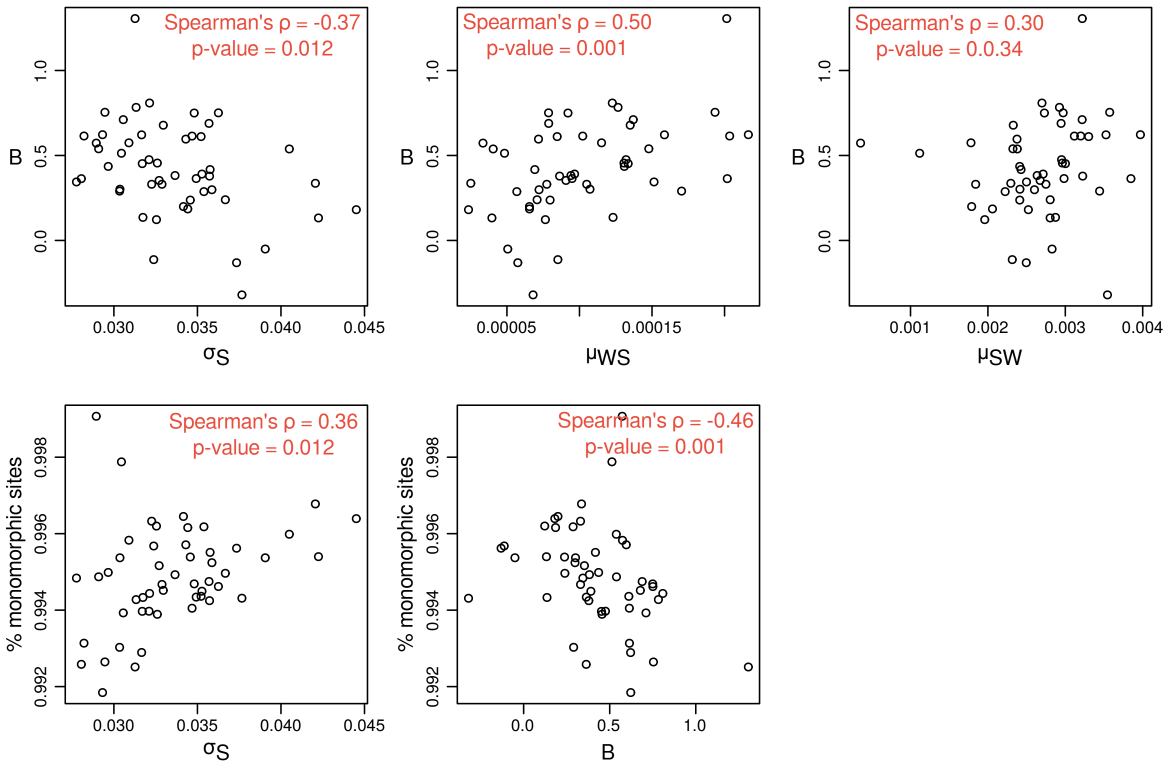


Figure S8: **Comparison of gBGC estimates between Glémin et al. (2015) and our method.** Estimates of gBGC rate coefficients were obtained using Glémin et al. (2015) and the method proposed here ($B$ in and $\sigma_S$, respectively). We used a human data set available as supplementary material in Glémin et al. (2015). The alleles counts correspond to 51 regions of 1 million sites from the human chromosome 1. We adapted our 4-variate model to only account for these two types of alleles: S and W stand for strong and weak alleles, respectively. The code needed to reproduce these analyses can be found in `https://github.com/pomo-dev/pomo_selection`. The correlation tests were FDR corrected for multiple testing.

| | Monomorphic | | | | Polymorphic | | | | | |
|---|---|---|---|---|---|---|---|---|---|---|
| **Population** | *A* | *C* | *G* | *T* | *AC* | *AG* | *AT* | *CG* | *CT* | *GT* |
| African humans | 615878 | 845533 | 666393 | 694569 | 249 | 1972 | 112 | 395 | 1716 | 318 |
| Non-african humans | 615966 | 845701 | 666455 | 694604 | 245 | 1780 | 101 | 321 | 1508 | 275 |
| Eastern gorillas | 615513 | 844903 | 666554 | 694492 | 126 | 977 | 60 | 192 | 874 | 139 |
| Western gorillas | 612727 | 839959 | 661019 | 691995 | 389 | 3238 | 180 | 506 | 2707 | 372 |
| Bonobos | 615217 | 844431 | 665216 | 693948 | 280 | 2422 | 134 | 408 | 1904 | 280 |
| Nigeria-Cameroon chimpanzees | 614656 | 843715 | 664742 | 693631 | 484 | 3888 | 227 | 632 | 3294 | 470 |
| Eastern chimpanzees | 614403 | 843458 | 664327 | 693337 | 377 | 3317 | 203 | 552 | 2593 | 409 |
| Central chimpanzees | 614292 | 843310 | 664183 | 693270 | 440 | 3369 | 214 | 521 | 2626 | 460 |
| Western chimpanzees | 615125 | 844759 | 665810 | 693893 | 246 | 1876 | 107 | 335 | 1499 | 261 |
| Sumatran orangutans | 615164 | 842548 | 663805 | 694268 | 414 | 3980 | 181 | 598 | 3192 | 468 |
| Bornean orangutans | 615573 | 843480 | 664912 | 694666 | 333 | 2655 | 135 | 420 | 2274 | 320 |

Table S1: **Summary of great apes' count data**.

| Scheme | $\pi_A$ | $\pi_C$ | $\pi_G$ | $\pi_T$ | $\rho_{AC}$ | $\rho_{AG}$ | $\rho_{AT}$ | $\rho_{CG}$ | $\rho_{CT}$ | $\rho_{GT}$ | $\sigma_A$ | $\sigma_C$ | $\sigma_G$ | $\sigma_T$ |
|---|---|---|---|---|---|---|---|---|---|---|---|---|---|---|
| M1 | 0.25 | 0.25 | 0.25 | 0.25 | 0.001 | 0.001 | 0.001 | 0.001 | 0.001 | 0.001 | - | - | - | - |
| M2 | 0.22 | 0.30 | 0.23 | 0.25 | 0.00028 | 0.00300 | 0.00016 | 0.00036 | 0.00172 | 0.00033 | - | - | - | - |
| S1 | 0.25 | 0.25 | 0.25 | 0.25 | 0.001 | 0.001 | 0.001 | 0.001 | 0.001 | 0.001 | 1 | 1 | 1 | 1 |
| S2 | 0.22 | 0.30 | 0.23 | 0.25 | 0.00028 | 0.00300 | 0.00016 | 0.00036 | 0.00172 | 0.00033 | 1 | 1.030 | 1.024 | 1.004 |

Table S2: **Simulation schemes.** Simulation schemes used to validate the Bayesian algorithms for estimating the model parameters under the multivariate Moran model with mutation (M schemes) and mutation plus selection (S schemes). $\sigma_A$ is set to 1.

| Population | $\mu_{AC}$ | $\mu_{CA}$ | $\mu_{AG}$ | $\mu_{GA}$ |
|---|---|---|---|---|
| African humans | 0.000237 | 0.000934 | 0.002248 | 0.008030 |
| Non-african humans | 0.000464 | 0.000957 | 0.003411 | 0.008700 |
| Eastern gorillas | 0.000220 | 0.000257 | 0.001850 | 0.002280 |
| Western gorillas | 0.001383 | 0.005259 | 0.014739 | 0.049773 |
| Bonobos | 0.000617 | 0.002196 | 0.005840 | 0.022977 |
| Nigeria-Cameroon chimpanzees | 0.000896 | 0.003185 | 0.008399 | 0.029962 |
| Eastern chimpanzees | 0.000516 | 0.001829 | 0.005393 | 0.018297 |
| Central chimpanzees | 0.000391 | 0.002039 | 0.003698 | 0.017267 |
| Western chimpanzees | 0.000391 | 0.000913 | 0.002932 | 0.008944 |
| Sumatran orangutans | 0.000573 | 0.001699 | 0.006940 | 0.017486 |
| Bornean orangutans | 0.000604 | 0.001116 | 0.005353 | 0.010287 |

| Population | $\mu_{AT}$ | $\mu_{TA}$ | $\mu_{CG}$ | $\mu_{GC}$ |
|---|---|---|---|---|
| African humans | 0.000224 | 0.000233 | 0.000982 | 0.000888 |
| Non-african humans | 0.000318 | 0.000296 | 0.000838 | 0.001036 |
| Eastern gorillas | 0.000114 | 0.000134 | 0.000352 | 0.000373 |
| Western gorillas | 0.001508 | 0.001765 | 0.004350 | 0.003859 |
| Bonobos | 0.000761 | 0.000639 | 0.001869 | 0.002065 |
| Nigeria-Cameroon chimpanzees | 0.000998 | 0.000964 | 0.002558 | 0.002566 |
| Eastern chimpanzees | 0.000594 | 0.000655 | 0.001700 | 0.001626 |
| Central chimpanzees | 0.000511 | 0.000514 | 0.001457 | 0.001304 |
| Western chimpanzees | 0.000289 | 0.000293 | 0.000788 | 0.001028 |
| Sumatran orangutans | 0.000475 | 0.000515 | 0.001739 | 0.001477 |
| Bornean orangutans | 0.000359 | 0.000378 | 0.001065 | 0.001108 |

| Population | $\mu_{CT}$ | $\mu_{TC}$ | $\mu_{GT}$ | $\mu_{TG}$ |
|---|---|---|---|---|
| African humans | 0.006177 | 0.001625 | 0.001235 | 0.000360 |
| Non-african humans | 0.005777 | 0.002607 | 0.001324 | 0.000484 |
| Eastern gorillas | 0.001609 | 0.001619 | 0.000291 | 0.000277 |
| Western gorillas | 0.033763 | 0.010382 | 0.005220 | 0.001810 |
| Bonobos | 0.015136 | 0.003569 | 0.002698 | 0.000576 |
| Nigeria-Cameroon chimpanzees | 0.021183 | 0.005752 | 0.003521 | 0.000954 |
| Eastern chimpanzees | 0.011799 | 0.003667 | 0.002102 | 0.000683 |
| Central chimpanzees | 0.011798 | 0.002272 | 0.002279 | 0.000491 |
| Western chimpanzees | 0.005329 | 0.002309 | 0.001194 | 0.000397 |
| Sumatran orangutans | 0.012329 | 0.004508 | 0.001913 | 0.000825 |
| Bornean orangutans | 0.007159 | 0.004084 | 0.001160 | 0.000636 |

| Population | $\sigma_C$ | $\sigma_G$ | $\sigma_T$ |
|---|---|---|---|
| African humans | 2.415 | 1.864 | 0.193 |
| Non-african humans | 1.192 | 1.161 | 0.053 |
| Eastern gorillas | 0.592 | 0.357 | 0.346 |
| Western gorillas | 1.711 | 1.334 | 0.285 |
| Bonobos | 1.705 | 1.551 | -0.057 |
| Nigeria-Cameroon chimpanzees | 1.741 | 1.473 | 0.091 |
| Eastern chimpanzees | 1.858 | 1.506 | 0.241 |
| Central chimpanzees | 2.600 | 2.083 | 0.145 |
| Western chimpanzees | 1.385 | 1.420 | 0.150 |
| Sumatran orangutans | 1.687 | 1.176 | 0.228 |
| Bornean orangutans | 1.087 | 0.846 | 0.194 |

Table S3: **Great apes mutation rates and selection coefficients.** Scaled mutation rates and selection coefficients estimated for the great apes populations using the multivariate Moran model with boundary mutations and allelic selection.
